# MEIS1 is Required for Establishing Bergmann Glia–Specific Properties in the Developing Cerebellum

**DOI:** 10.64898/2026.01.21.700745

**Authors:** Kentaro Ichijo, Toma Adachi, Tomoo Owa, Minami Mizuno, Kyoka Suyama, Kaiyuan Ji, Koichi Hashizume, Ikuko Hasegawa, Eriko Isogai, Masaki Sone, Yukiko U. Inoue, Ryo Goitsuka, Takuro Nakamura, Takayoshi Inoue, Satoshi Miyashita, Kenji Kondo, Tatsuya Yamasoba, Mikio Hoshino

## Abstract

Compared to normal multipolar astrocytes, Bergmann glial cells (BGs), specifically differentiated astrocytes in the cerebellum, possess unique unipolar morphology and additional cellular functions. However, the molecular mechanisms that confer BG-specific properties onto normal multipolar astrocytes remain unknown. Here, we show that the transcription factor, MEIS1, is involved in BGs acquiring their unique characteristics. Targeted disruption of *Meis1* in the whole cerebellum or astroglial lineage cells resulted in a marked reduction of BGs accompanied by an increase in multipolar astrocytes in mice. Postnatal deletion of *Meis1* in Bergmann glia-like progenitors (BGLPs), which produce both BGs and multipolar astrocytes, suppressed their differentiation into BGs while promoting into multipolar astrocytes. Single-cell RNA sequencing, immunohistochemistry, and ChIP-Atlas analyses indicated that MEIS1 directly upregulates expression of BG-specific genes, including *Vimentin* and *Zeb2*, which are known to contribute to the correct localization and the unipolar process formation of BGs. These findings suggest that MEIS1 promotes the endowment of BG-specific properties to astrocytes by controlling the expression of BG-specific genes, thereby ensuring proper differentiation of BGLPs into BGs.

## Introduction

Astrocytes are widely distributed throughout the central nervous system and play multiple essential roles, including providing structural support for synapses, regulating metabolism, uptaking neurotransmitters, maintaining ionic homeostasis, and contributing to the formation of the blood–brain barrier (Khakh and Sofroniew, 2015; Sofroniew and Vinters, 2010). In the mature mammalian cerebellum, astrocytes are localized in three distinct regions. In the inner granule cell layer (IGL) and white matter (WM), astrocytes with a typical multipolar morphology commonly observed in the mammalian brain are present. In contrast, in the Purkinje cell layer (PCL), the cell body of Bergmann glial cells (BGs), a specialized type of astrocyte with a characteristic unipolar morphology, are aligned in a single row, extending their long processes radially through the molecular layer (ML) (Leto et al., 2016). BGs not only share the basic properties of astrocytes but also possess distinct features: during development, their elongated radial fibers serve as scaffolds for granule cell migration (Xu et al., 2013), and in the mature cerebellum, they contribute to the maintenance of cortical architecture (Hull et al., 2023). Thus, BGs are specialized astrocytes endowed with additional morphological and functional characteristics; however, the detailed molecular basis that confers these unique properties remains unknown. (In this study, we collectively refer to BGs and multipolar astrocytes as “astroglial cells,” while the term “astrocytes” specifically denotes the multipolar type.) In the postnatal cerebellum of mammals, astroglial cells are generated from two distinct types of progenitors (astroglial progenitors). One is an astrocyte-like progenitor (AsLP) localized in the WM with a multipolar morphology, and the other is a Bergmann glia-like progenitor (BGLP) that exhibits a unipolar morphology and a localization pattern similar to that of BG. AsLPs primarily give rise to astrocytes (Permigiani et al., 2015), whereas BGLPs produce both BGs and astrocytes (Suyama et al., 2025, bioRxiv; Cerrato et al., 2018). However, the molecular mechanisms that determine how BGLPs generate each astroglial subtype remain poorly understood.

MEIS1 is a transcription factor belonging to the three amino acid loop extension (TALE) homeodomain protein family (Hisa et al., 2004; Azcoitia et al., 2005). In the embryonic cerebellar primordium of mice, MEIS1 is expressed in the neuroepithelial cells of the fourth ventricular zone (VZ). In the postnatal cerebellum, its expression is observed in granule cells (GCs), their progenitors (granule cell progenitors (GCPs)), and at least a subset of SOX9-positive astroglial lineage cells (astroglial progenitors and astroglial cells) (Owa et al., 2018). We have previously shown that MEIS1 regulates the differentiation of GCPs into GCs by controlling *Pax6* transcription, BMP signaling, and ATOH1 degradation (Owa et al., 2018). However, the role of MEIS1 in non-granule cell lineages of the cerebellum has remained unexplored.

In this study, we first found that cerebellum-wide deletion of *Meis1* caused cerebellar hypoplasia and motor learning deficits, together with a marked reduction of BGs and an increase in multipolar astrocytes. Moreover, astroglial lineage–specific deletion of *Meis1* also resulted in decreased production of BGs from BGLPs and a concomitant increase in astrocytes, suggesting that MEIS1 contributes to the differentiation of BGLPs into BGs. Single-cell RNA sequencing, immunohistochemistry, and ChIP-Atlas analyses further suggested that MEIS1 promotes the expression of BG-like cell–specific genes, including *Vimentin* and *Zeb2*. These genes have been reported to contribute to the formation or maintenance of the characteristic unipolar radial processes of BGs, and the reduced expression of these genes in *Meis1*-deficient mice is consistent with the disruption of BG-specific properties. Taken together, these findings suggest that MEIS1 confers BG-specific characteristics on postnatal cerebellar astrocytes by regulating the expression of BG-like cell–specific genes.

## Results

### *Meis1* deficiency throughout the cerebellum causes severe impairments in cerebellar structure and function

During cerebellar development, MEIS1 is expressed in PAX6-expressing cells (GCPs and GCs) and SOX9-expressing cells (astroglial lineage cells) as well as in radial glial cells in the VZ (Owa et al., 2018). Using conditional knockout (cKO) mice specific for GC-lineage cells (*Meis1^fl/fl^; Atoh1-Cre-Tg*), we previously showed that MEIS1 has a cell-autonomous function to promote the degradation of ATOH1 in GCPs and thereby enhancing their differentiation into GCs (Owa et al., 2018). By crossing *Meis1 ^fl^* and *En1^Cre^* (Kimmel et al., 2000) allele-carrying mice, we generated *Meis1^fl/fl^; En1^Cre/+^* mice, in which the *Meis1* gene product was deleted in the entire cerebellum during development (Figure S1a, b). In this study, these two types of cKO mice are referred to as *Atoh1::Meis1cKO* and *En1::Meis1cKO*, respectively.

At P21, *En1::Meis1* cKO mice exhibited more severe cerebellar hypoplasia (Figure 1a) than previously reported *Atoh1::Meis1cKO* mice (Owa et al., 2018). Furthermore, *En1::Meis1* cKO cerebellum lacked the vermis (Figure 1a). At two months of age, the rotarod test revealed obvious motor learning impairments in *En1::Meis1cKO* mice, but no such impairments were observed in *Atoh1::Meis1cKO* mice (Figure 1b). Nissl staining highlighted that lobular structure and layer structure were mildly and severely disrupted in *Atoh1::Meis1cKO* and *En1::Meis1cKO*, respectively (Figure S2a).

**Figure 1.**
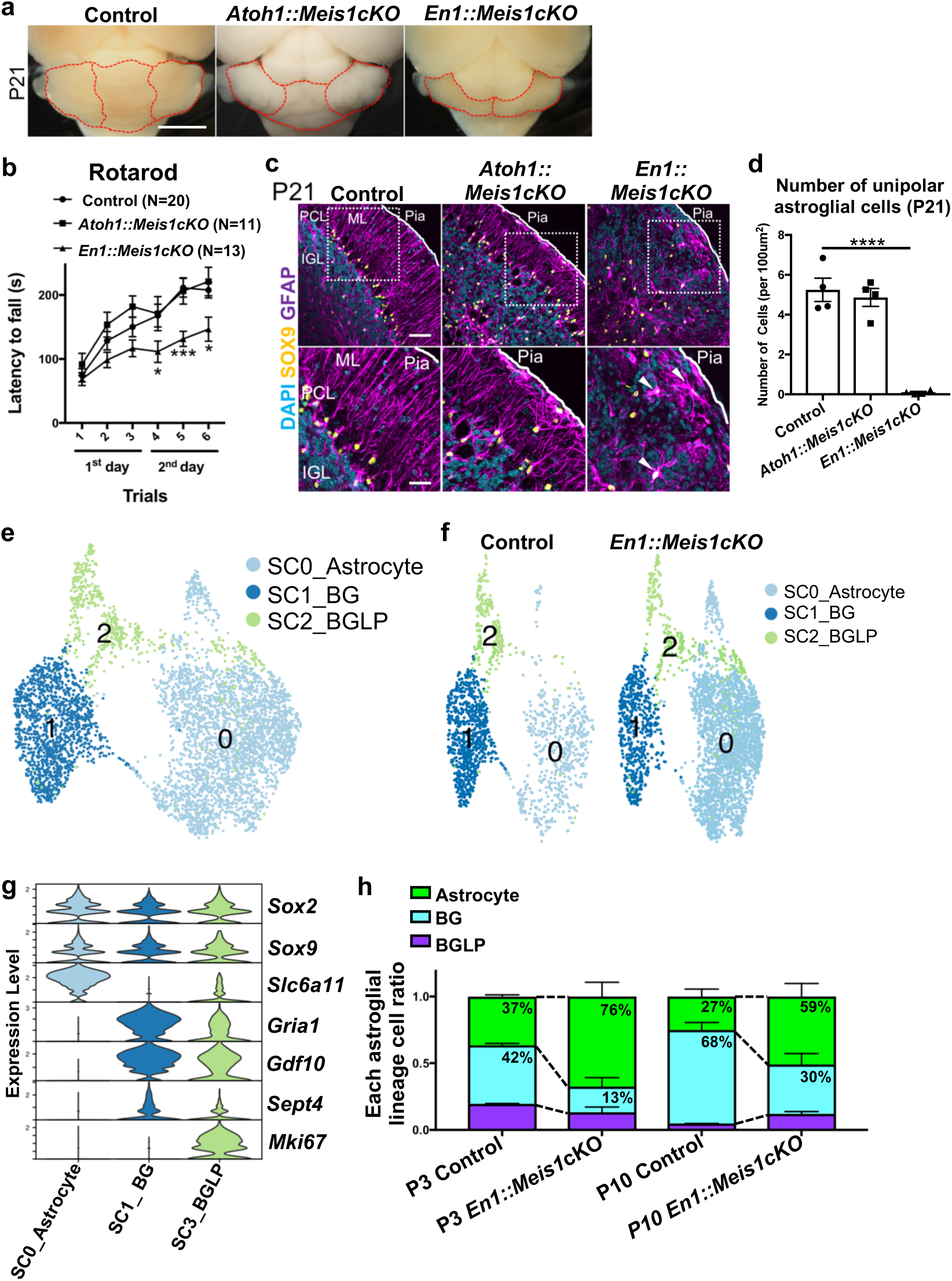
Loss of *Meis1* in the whole cerebellum disrupts cerebellar architecture, impairs motor learning, and reduces BG-like cells (a) Bright-field images of cerebella from control, *Atoh1::Meis1cKO*, and *En1::Meis1cKO* at P21. Dashed lines indicate the boundaries between the cerebellar hemisphere and vermis. Scale bar, 3 mm. (b) Latency to fall in the rotarod test for control (N = 20), *Atoh1::Meis1cKO* (N = 11), and *En1::Meis1cKO* (N = 13) at 2 months of age. Statistical tests were performed by Tukey’s multiple comparisons test; p values were represented as; *** for p < 0.001, * for p < 0.05. (c) Immunostaining of P21 cerebellar sections from each genotype with DAPI, SOX9, and GFAP. White arrowheads indicate ectopically localized multipolar astrocytes near the cerebellar surface. Scale bars: 50 µm (upper panels), 25 µm (lower panels). (d) Quantification of the unipolar astroglial cell (BG-like cell) number in the P21 cerebellum of each genotype. Statistical analysis was performed using Dunnett’s multiple comparisons test. **** indicates p < 0.0001. (e, f) Uniform manifold approximation and projection (UMAP) dimensional reduction of astroglial lineage cell cluster (Cluster 4, 7,504 cells). Three distinct sub-clusters were identified (e), and their distribution is shown separately for Control and *En1::Meis1cKO* cerebella (f). (g) Violin plots showing the expression of proliferation marker (*Mki67*), BG-like cell markers (*Gria1, Gdf10, Sept4*), and markers highly expressed in astrocytes (*Slc6a11*) across the three sub-clusters. (h) Stacked bar graphs showing the proportion of each astroglial lineage cell type at P3 and P10 in control and *En1::Meis1cKO* cerebella.

In *En1::Meis1cKO*, the degree of layer structure defects was stronger in the rostral side and weaker in the caudal side (Figure S2, S3). The severe impaired structure was also evident at earlier stages, such as P3 and P10, in *En1::Meis1cKO* cerebella (Figure S4a). Immunohistochemical staining for cleaved caspase-3 showed no increase in total apoptotic cells or astroglial lineage (SOX9-positive) apoptotic cells in *En1::Meis1* cKO cerebella at either P3 or P10 compared to control (Figure S5). This suggests that the disorder in the cerebellar structure of *En1::Meis1* cKO was not caused by cell death, but rather by some kind of developmental abnormality.

As previously reported (Owa et al., 2018), in the *Atoh1::Meis1cKO* cerebella at P21, there were no major disruptions in the arrangement of Purkinje cells (CALBINDIN-positive cells, PCs,) or GCs (NEUROD1-positive cells), despite minor cellular dislocations (Figure S2b, c). However, in *En1::Meis1cKO* at P21, the arrangement of both types of cells was significantly destroyed (Figure S2b, c). In particular, GCs remained in the cerebellar surface comprising the external granule cell layer (EGL)-like structure and IGL was hardly formed especially in the rostral side (Figure S2a, c).

Similarly, dislocation of PCs was also observed at earlier stages, especially at P10 (Figure S4b).

### Astroglial lineage cells in *En1::Meis1cKO* cerebella

The phenotypic divergence between the two types of *Meis1* cKO mice became particularly apparent after birth (see below) and was likely due to the loss of MEIS1 expression in SOX9-positive cells (astroglial lineage cells) in addition to the GC lineage cells in *En1::Meis1cKO* cerebella. To investigate which astroglial lineage cells express MEIS1 in the cerebellum after birth, we performed immunostaining on P0 and P10 cerebella with KI67 (a marker of cell proliferation), and SOX9 and VIMENTIN (for P0) or GFAP (for P10), markers of astroglial lineage cells (Figure S6). MEIS1 protein expression was observed in all astroglial lineage cells, namely BGLPs, AsLPs, BGs, and astrocytes (Figure S6). However, as was reported previously, MEIS1 expression was lost until P21, when the cerebellar structure is established (Owa et al., 2018).

To investigate the phenotype of astroglial lineage cells after establishment of the cerebellar structure, we performed immunostaining using SOX9 and GFAP, which are their nuclear and cytoplasmic markers, respectively, on control (*Meis1^fl/fl^*) and the two types of cKO cerebella at P21 (Figure 1c). First, we focused on the rostral cerebellum, where more severe structural impairments were observed in *En1::Meis1* cKO. In control cerebellum, the cell bodies of BGs lined up with the PCL, and their unipolar projections extended toward the pia mater. In *Atoh1::Meis1cKO*, most BG cell bodies were localized in the PCL and still had long unipolar projections toward the pia, although a small population of BG cell bodies were ectopically located in the ML (Figure 1c). However, astroglial (SOX9-positive) cells with a unipolar structure like BGs were rarely observed in *En1::Meis1* cKO (Figure 1c, d). Interestingly, astroglial (SOX9-positive) cells with multipolar processes, which resembled the morphology of astrocytes, were observed near the surface of the *En1::Meis1* cKO cerebellum (Figure 1c, arrowheads). However, as expected, in the caudal side of *En1::Meis1* cKO at P21, these phenotypes were quite mild (Figure S3c), and there was not significant change in the number of astroglial (SOX9-positive) cells with a unipolar structure like BGs (Figure S3d). Consistently, the misalignment of PCs and GCs at the caudal side of P21 *En1::Meis1* cKO was quite minor compared to the rostral side (Figure S3a, b).

Immunostaining showed that the decrease in BGs (or unipolar astroglial cells) and the appearance of ectopic multipolar astroglial cells in the cerebellar surface were also observed at earlier postnatal stages, such as P3 and P10, in *En1::Meis1* cKO (Figure S7). However, at an embryonic stage, E15.5, the number of astroglial lineage cells was not significantly changed in *En1::Meis1* cKO cerebella (Figure S8). Because these astroglial lineage cells at embryonic stages are derived from radial glial cells in the VZ (Buffo et al., 2013), it was suggested that loss of *Meis1* expression in the VZ did not seem to affect the production of astroglial lineage cells.

### Single cell RNA-sequencing analysis of *En1::Meis1cKO*

To evaluate changes in gene expression characteristics of astroglial lineage cells due to the deficiency of *Meis1*, we performed single cell RNA-sequencing (scRNA-seq) on control and *En1::Meis1* cKO at P3 and P10 (Figure 1e-h, Figure S9). Two individual cerebella from each group were used. In total, we successfully obtained data from 79,038 cells. Using unsupervised clustering, we obtained 17 clusters shared by the two genotypes (Figure S9a-e). Each cluster was annotated based on the expression of known marker genes corresponding to distinct cerebellar cell types (Figure S9f). During this annotation process, clusters 4 showed high expression of astroglial lineage markers (*Aldoc, Aqp4, Sox2, Sox9, S100b, Fgfr1, Fabp7, Hopx*) (Figure S9f, g). For further analysis, unsupervised subclustering was performed on cluster 4 (7,504 cells), and three subclusters were identified (Figure 1e, f). Annotation for each subcluster using proliferation marker (*Mki67*), BG-like cell (BGLPs and BGs) specific markers (*Gria1, Gdf10, Sept4*) (Saab et al., 2012; Mecklenburg et al., 2014; Ageta-Ishihara et al., 2013), and markers highly expressed in astrocyte-like cells (AsLPs, and astrocytes) (*Slc6a11*) (Bayin et al., 2021) suggested that subclusters 0 corresponded to astrocytes, subcluster 2 corresponded to BGLPs and subcluster 1 corresponded to BGs (Figure 1e-g). Based on the number of cells in each subcluster, it was suggested that the proportions of BGLPs and BGs were clearly reduced in *En1::Meis1*cKO cerebella, while that of astrocytes increased (Figure 1h). This is consistent with the reduction of unipolar astroglial cells in *En1::Meis1*cKO cerebella observed in histological analysis (Figure 1c, d, Figure S7). This observation may imply that the ectopic multipolar shaped astroglial cells observed in the *En1::Meis1*cKO cerebella (Figure 1c, d, Figure S7) were cells that should have become BGs but lost their BG-specific properties and became normal astrocytes. In other words, it is suggested that MEIS1 confers additional properties to normal astrocytes and turns them into BGs.

### Deletion of *Meis1* gene in cultured astroglial lineage cells

Previously, Martínez-Lozada et al. developed an experimental system for culturing solely astroglial lineage cells in the cerebellum (Martínez-Lozada et al., 2011). Tellios et al. reported that, in this culture system, after the cell passage, approximately 90% of the cells are unipolar BG-like cells, whereas the remaining cells exhibit multipolar astrocyte-like morphology (Tellios et al., 2021; Martínez-Lozada et al., 2011). Using the same method, we isolated and cultured astroglial lineage cells from cerebella of three genotypes, WT, *Meis1^fl/fl^,* and *Meis1^fl/fl^; Sox9^CreERT2/+^* (Figure S10). At 3 days or 7 days after passage, immunostaining was performed with GFAP, GLAST, KI67 and MEIS1 (Figure S10a-d). Based on cell morphology and immunoreactivity, we differentially identified BGLPs (unipolar, KI67-positive), BGs (unipolar, KI67-negative), AsLPs (multipolar, KI67-positive). In our experimental conditions, astrocytes (multipolar, KI67- negative) were not observed. The ratio of cell types did not change among genotypes either at 3 days or 7 days after passage (Figure S10e, f).

*Sox9^CreERT2/+^* allele is designed to express CreERT2 protein in SOX9-positive cells, in which Cre-recombinase is activated by administration of tamoxifen or its derivatives (Soeda et al., 2010.). At three days after passage, we administrated 5 or 10 µM of 4-hydroxytamoxifen (4-OH-TX) to cultured astroglial cells from *Meis1^fl/fl^,* or *Meis1^fl/fl^; Sox9^CreERT2/+^* cerebellum (Figure 2a). Cells were harvested at seven days after passage and subjected to quantitative reverse transcription polymerase chain reaction (qRT-PCR) using primer pairs for *Meis1* and *Sox2*. At either concentration of 4-OH-TX, a significant decrease in *Meis1* expression was observed in *Meis1^fl/fl^; Sox9^CreERT2/+^* cells but not in control *Meis1^fl/fl^* cells, as expected (Figure 2b). In contrast, *Sox2* expression levels did not differ under any conditions (Figure 2b). These observations suggest that the conditional deletion experiment of *Meis1* induced by 4-OH-TX functions appropriately.

**Figure 2.**
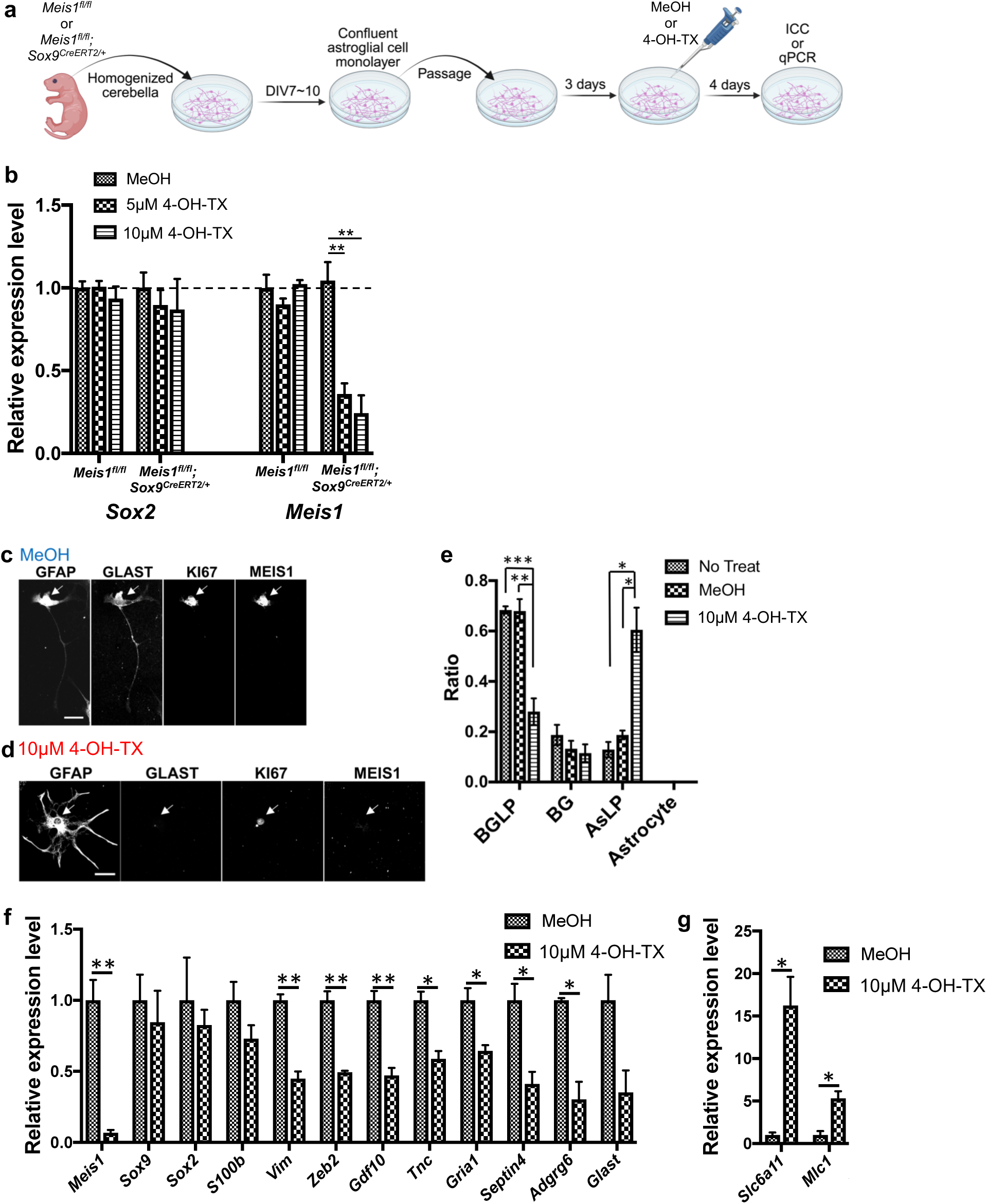
*Meis1* knockout decreases BG-like cells and increases astrocyte-like cells in cerebellar astroglial cell culture (a) Experimental scheme. Cerebella were isolated from P0–P2 *Meis1^fl/fl^*and *Meis1^fl/fl^; Sox9^CreERT2/+^* mice, dissociated, and cultured. When cultures reached confluence at DIV7–10, cells were passaged at a density of 1×10^5^ cells per well. Three days later, either MeOH (control) or 4-hydroxytamoxifen (4-OH-TX) was added. Four days after treatment, immunocytochemistry (ICC) or quantitative PCR (qPCR) was performed. (b) qPCR analysis of *Sox2* and *Meis1* expression in *Meis1^fl/fl^*and *Meis1^fl/fl^; Sox9^CreERT2/+^* cultures treated with MeOH (control), 5 µM 4-OH-TX, or 10 µM 4-OH-TX. Data are shown as relative expression levels normalized to control (MeOH treated *Meis1^fl/fl^*). Statistical analysis was performed using Dunnett’s multiple comparisons test; ** indicates p < 0.01. (c, d) Immunocytochemistry of cultured cerebellar astroglial cells with GFAP, GLAST, KI67, and MEIS1. GLAST expression in cerebellar astroglial cell culture is known to be restricted to BG-like cells (Tellios et al., 2021). Representative images show the predominant cell type in *Meis1^fl/fl^; Sox9^CreERT2/+^*derived astroglial cultures treated with MeOH (c) and with 10 µM 4-OH-TX (d). Scale bars,10 µm. (e) Ratios of astroglial lineage cell types in cultured astroglial cells derived from *Meis1^fl/fl^; Sox9^CreERT2/+^* mice under three conditions: no treatment, MeOH, and 10 µM 4-OH-TX. Statistical analysis was performed using Dunnett’s multiple comparisons test. p values were represented as * for p < 0.05, ** for p < 0.01, and *** for p < 0.001. (f, g) qPCR analysis of astroglial cell cultures derived from *Meis1^fl/fl^; Sox9^CreERT2/+^* mice treated with MeOH or 10 µM 4-OH-TX. Relative expression levels of *Meis1*, astroglial cell common markers (*Sox9*, *Sox2*, *S100b*) and BG-like cell–specific markers (*Vim*, *Zeb2*, *Gdf10*, *Tnc*, *Gria1*, *Septin4*, *Adgrg6*, *Glast*) were shown in (f). Relative expression levels of *Slc6a11,* which is known to be highly expressed in astrocyte-like cells, were shown in (g). Statistical analysis was performed using two-tailed t-tests; p values were indicated as ** for p < 0.01, * for p < 0.05.

Next, we cultured astroglial cells from *Meis1^fl/fl^; Sox9^CreERT2/+^* cerebellum and subsequently added control solvent or 4-OH-TX (10 µM) at three days after passage. At seven days after passage, cells were subjected to immunostaining with specific antibodies or qRT-PCR with specific primer pairs. Compared to control, MEIS1 protein signal was under detection level in immunostaining and *Meis1* transcript level was much reduced (Figure 2c, d, f), suggesting that conditional *Meis1* deletion was successfully achieved, as expected. Identification of cell types based on their morphologies and immunoreactivities revealed that conditional deletion of *Meis1* resulted in a significant decrease in BGLPs and a dramatic increase in AsLPs (Figure 2c-e). The qRT-PCR experiment showed that the *Meis1* deletion did not affect the expression of markers common to astroglial lineage cells (*Sox9, Sox2, S100b*).

However, genes specific to BG-like cells (BGLPs, BGs) (*Vim, Zeb2, Gdf10, Tnc, Gria1, Septin4, Adgrg6*) (Ribotta et al.,2000; He et al., 2018; Mecklenburg et al., 2014; Bartsch et al., 1992; Saab et al., 2012; Ageta-Ishihara et al., 2013; Koirara et al., 2010) all showed a significant reduction in expression by *Meis1* deletion (Figure 2f). In contrast, *Slc6a11*, a gene known to be highly expressed in astrocyte-like cells (AsLPs, astrocytes) (Bayin et al., 2021), showed increased expression in those *Meis1*-deleted cells (Figure 2g) These *in vitro* observations further suggest that MEIS1 protein, expressed in astroglial linage cells, is involved in their differentiation into the specialized astrocytes, or BGs.

### Conditional deletion of *Meis1* in astroglial lineage cells after birth

We next tried to generate the cKO mice in which the *Meis1* gene was deleted specifically in astroglial lineage cells after birth. *Meis1^fl/fl^*or *Meis1^fl/fl^; Sox9^CreERT2/+^* mice were administrated with tamoxifen via intraperitoneal injection at P0 and P1, and subjected to immunohistochemical analysis at P10 (Figure 3a). We have reported that astroglial progenitors (BGLPs, AsLPs) disappear as early as P10 during cerebellar development (Suyama et al, 2025. bioRxiv). In the *Meis1^fl/fl^; Sox9^CreERT2/+^* cerebella, MEIS1 signal intensities were much reduced in astroglial lineage (SOX9-positive) cells, but not in cells in the EGL, compared to control (Figure S11a-c), confirming that *Meis1* gene was specifically deleted in astroglial lineage cells in this experimental condition.

**Figure 3.**
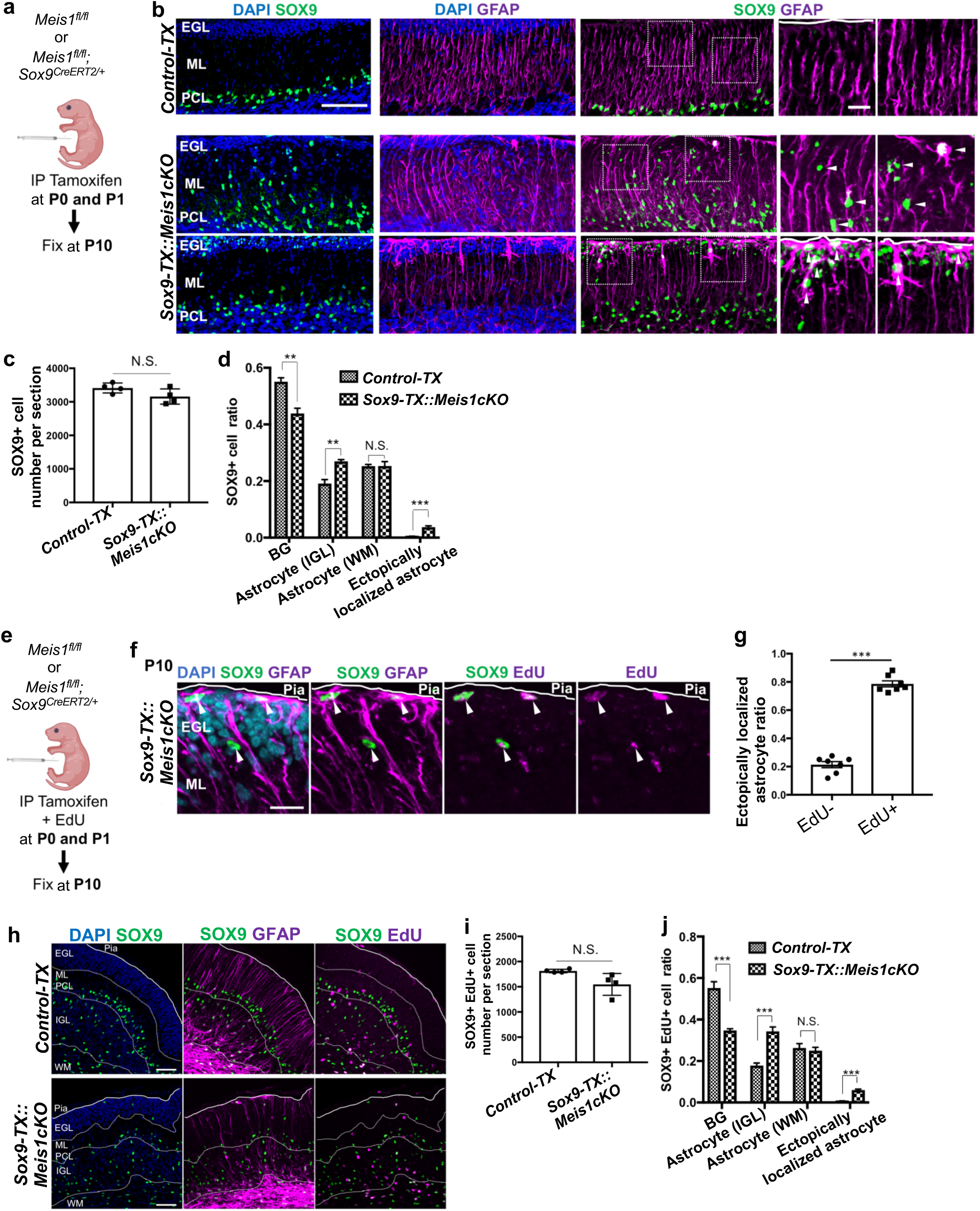
Astroglial lineage-specific knockout of *Meis1* in postnatal stage leads to a decrease in BGs and an increase in astrocytes (a) Experimental scheme. Tamoxifen was intraperitoneally (IP) injected to *Meis1^fl/fl^*or *Meis1^fl/fl^; Sox9^CreERT2/+^* mice at P0 and P1, and the cerebella were fixed at P10 for subsequent analyses. (b) Immunostaining of P10 cerebellar sections from *Control-TX* and *Sox9-TX::Meis1cKO* with SOX9 and GFAP. White arrowheads indicate ectopically localized multipolar astrocytes in the molecular layer (ML) and external granule cell layer (EGL). Scale bars: 50 µm (lower magnification), 25 µm (higher magnification). (c) Quantification of SOX9-positive cells in P10 cerebellar sections from *Control-TX* and *Sox9-TX::Meis1cKO*. Statistical analysis was performed using two-tailed t-tests. N.S. indicates p > 0.05. (d) Quantification of the proportions of SOX9-positive cells among different astroglial lineage cell types in P10 cerebellar sections from *Control-TX* and *Sox9-TX::Meis1cKO*. Statistical analysis was performed using paired t-tests. p values are represented as *** for p < 0.001, ** for p < 0.01, and N.S. for p > 0.05. (e) Experimental scheme. Tamoxifen and EdU were intraperitoneally (IP) injected to *Meis1^fl/fl^* or *Meis1^fl/fl^; Sox9^CreERT2/+^* mice at P0 and P1, and the cerebella were fixed at P10 for subsequent analyses. (f) Immunostaining of P10 cerebellar sections from *Sox9-TX::Meis1cKO* with SOX9, GFAP and EdU. White arrowheads indicate ectopically localized multipolar astrocytes as in (b). Scale bars: 25 µm. (g) Quantification of the proportion of ectopically localized astrocytes that were EdU-positive or EdU-negative in *Sox9-TX::Meis1cKO*. Statistical analysis was performed using two-tailed t-tests. p values are represented as *** for p < 0.001. (h) Immunostaining of P10 cerebellar sections from *Control-TX* and *Sox9-TX::Meis1cKO* with SOX9, GFAP and EdU. Scale bars: 50 µm. (i) Quantification of SOX9 and EdU double positive cells in P10 cerebellar sections from *Control-TX* and *Sox9-TX::Meis1cKO*. Statistical analysis was performed using two-tailed t-tests. N.S. indicates p > 0.05. (j) Quantification of the proportions of SOX9 and EdU double positive cells among different astroglial cell types in P10 cerebellar sections from *Control-TX* and *Sox9-TX::Meis1cKO*. Statistical analysis was performed using two-tailed t-tests. p values are represented as *** for p < 0.001 and N.S. for p > 0.05.

In this study, we call tamoxifen-administrated *Meis1^fl/fl^* and *Meis1^fl/fl^; Sox9^CreERT2/+^* mice as *control-TX* and *Sox9-TX:: Meis1cKO*, respectively.

For histological analyses, we performed immunostaining with SOX9 and GFAP to the cerebella of *control-TX* and *Sox9-TX:: Meis1cKO* mice at P10. No significant difference was observed in the total number of astroglial linage (SOX9-positive) cells (Figure 3b, c). However, BGs were significantly reduced and IGL astrocytes increased in *Sox9-TX:: Meis1cKO* cerebella, compared to *control-TX* (Figure 3b, d). Furthermore, ectopically localized astrocytes, observed in *En1:: Meis1cKO*, were also detected in the ML and EGL of *Sox9-TX:: Meis1cKO* cerebella (Figure 3b, d, arrowheads).

In *Sox9-TX:: Meis1cKO* cerebella, MEIS1 expression is lost in both mitotic cells (BGLPs, AsLPs) and postmitotic cells (BGs, astrocytes) in the astroglial lineage.

Therefore, we designed an experiment to distinguish whether the loss of *Meis1* in mitotic or postmitotic astroglial lineage cells is involved in the phenotype observed in *Sox9-TX:: Meis1cKO* cerebella. *Meis1^fl/fl^*or *Meis1^fl/fl^; Sox9^CreERT2/+^* mice were administrated with tamoxifen and EdU at P0 and P1, and subjected to immunohistochemical analysis at P10 (Figure 3e). In this experiment, EdU-positive cells at P10 are presumed to originate from cells that were mitotic at the timing of tamoxifen administration. However, since EdU is only incorporated into S phase cells at the timing of administration, we cannot conclude that EdU-negative cells at P10 are derived from postmitotic cells. Some of them might be derived from mitotic cells in G1, G2 or M phase when tamoxifen was administrated. Immunohistochemical analysis revealed that around 80% of ectopically localized astrocytes in *Sox9-TX:: Meis1cKO* cerebella were EdU-positive (Figure 3f, g), suggesting that most (at least 80%) of them originated from mitotic astroglial cells, namely, BLBPs or AsLPs. We found that cell numbers of EdU-positive astroglial (SOX9-positive) cells were not different between *control-TX* and *Sox9-TX:: Meis1cKO* (Figure 3h, i). However, among EdU-positive cells, the proportion of BGs decreased significantly, and those of astrocytes and ectopically localized astrocytes increased in *Sox9-TX:: Meis1cKO* compared to *control-TX* (Figure 3h, j). Interestingly, the changes in the proportion of astroglial lineage cells in *Sox9-TX:: Meis1cKO* cerebella were more pronounced when focusing on EdU-positive cells rather than all astroglial cells (Figure 3d, j). These observations suggest that majority of phenotypes observed in *Sox9-TX:: Meis1cKO* cerebella was caused by the loss of *Meis1* in astroglial progenitors (BGLPs, AsLPs) at the early postnatal stage.

### Conditional deletion of *Meis1* in BGLPs

We and other research groups have previously showed that BGLPs produce not only BGs but also astrocytes during postnatal cerebellar development (Suyama et al, 2025. bioRxiv; Cerrato et al., 2018). In contrast, AsLPs have been reported to only produce astrocytes, but not BGs (Parmigiani et al., 2015). These past findings led us to the notion that deficiency of *Meis1* in BGLPs may account for the imbalance in the differentiation into BGs and astrocytes, observed in *Sox9-TX:: Meis1cKO* cerebella (Figure 3d, j).

We previously developed a method to label BGLPs and their descendant cells with GFP by electroporating a human GFAP-promoter-driven EGFP (hGFAP-EGFP) vector into the cerebellar surface at early postnatal stages (Suyama et al, 2025. bioRxiv). We utilized hGFAP-EGFP-T2A-iCre vector, which was designed to express improved CRE (iCRE) recombinase in the same cells expressing EGFP through the action of the T2A peptide. By using this gene-transfer method, WT or *Meis1^fl/fl^*cerebella were electroporated with hGFAP-EGFP-T2A-iCre at P0 and subjected to further analyses at P10 (Figure 4). Immunostaining revealed that MEIS1 protein levels were under detection level in astroglial lineage (SOX9-positive) cells in *Meis1^fl/fl^* cerebella but not in control (WT) (Figure 4b), suggesting that the specific cKO for *Meis1* in BGLPs was successfully achieved. Observation of cell morphology and immunoreactivities suggested that the proportion of BGs decreased significantly, and that of astrocytes increased in *Meis1^fl/fl^* cerebella introduced with hGFAP-EGFP-T2A-iCre, compared to control (Figure 4c, d). These findings suggest that *Meis1* expression in BGLP is required for BG production from BGLP, and that loss of *Meis1* expression causes cells produced from BGLP to shift from BGs to astrocytes. In the cKO experiment, ectopically localized astrocytes were not observed. In these electroporation-based experiments, only a relatively small number of cells could be labeled with EGFP, so those ectopic astrocytes may have been missed in our observations.

**Figure 4.**
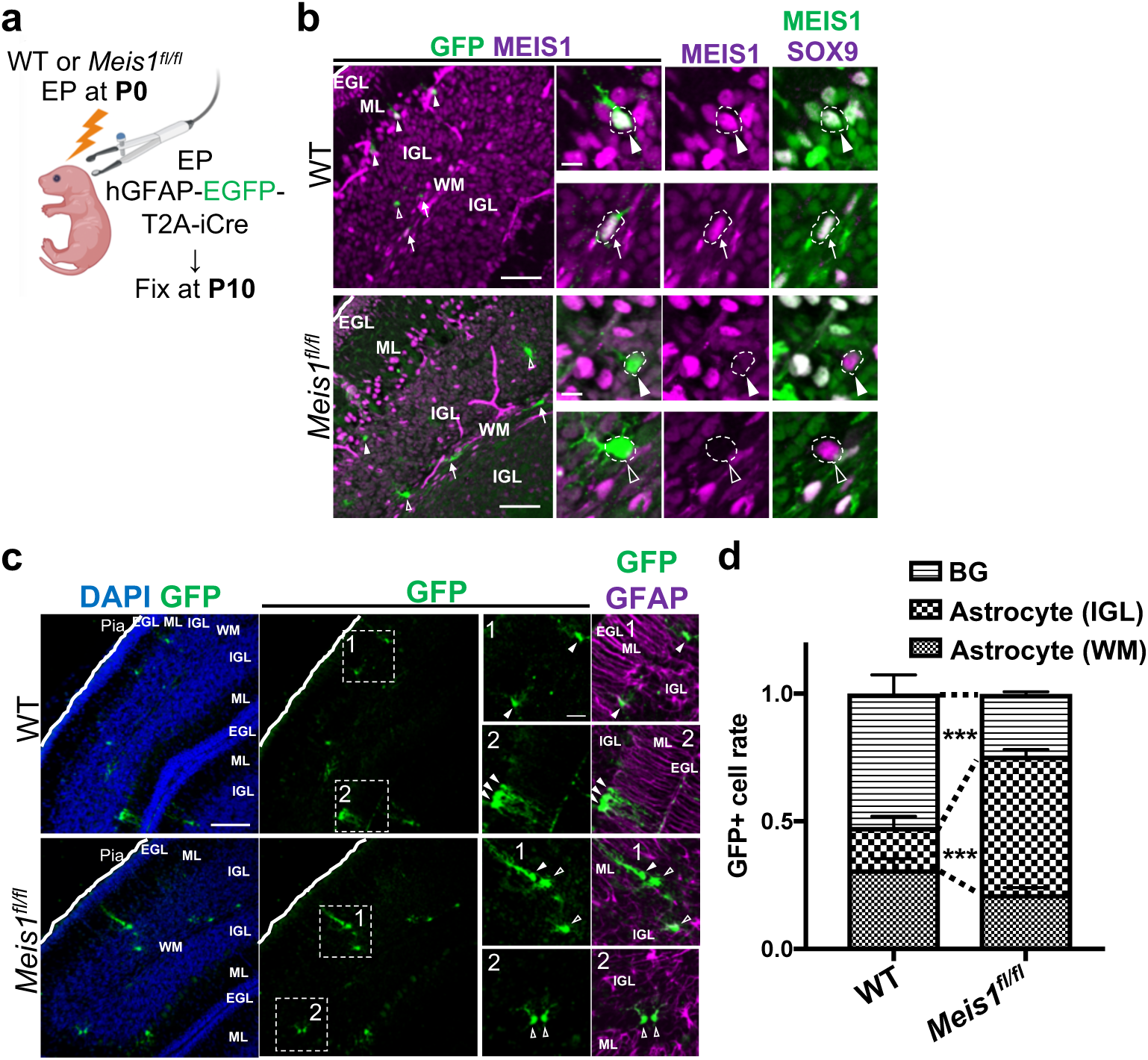
Loss of *Meis1* expression in BGLPs shifts the differentiation balance from BGs to astrocytes (a) Experimental scheme. Wild-type (WT) or *Meis1^fl/fl^* mice at P0 were electroporated (EP) with the hGFAP-EGFP-T2A-iCre vector into the cerebellar surface. The cerebella were fixed at P10 for subsequent analyses. (b, c) Immunostaining of P10 cerebellar sections from WT and *Meis1^fl/fl^* mice with GFP, SOX9 and MEIS1 (b), or GFP and GFAP (c). White arrowheads indicate GFP-positive BGs, white arrows indicate astrocytes in IGL, and open arrowheads (white-outlined) indicate astrocytes in WM. Scale bars: 100 µm (lower magnification of (b)), 20 µm (higher magnification of (b)), 100 µm (lower magnification of (c)), and 25 µm (higher magnification of (c)). (d) Quantification of GFP-positive astroglial cells in P10 cerebellar sections from WT and *Meis1^fl/fl^*mice. The proportions of GFP-positive BGs, IGL astrocytes, and WM astrocytes were compared between genotypes. Statistical analysis was performed using two-tailed t-tests; *** indicates p < 0.001.

### Downstream candidate genes of MEIS1 in BG-like cells

Since MEIS1 is a transcriptional activator, this protein is thought to regulate the expression of downstream genes in BG-like cells (BGLPs and BGs) for their proper characterization. To identify downstream target genes of MEIS1, we selected subclusters 1 and 2 of scRNA-seq data (Figure 1e) corresponding to BG-like cells and obtained 696 genes that showed significantly reduced expression in *En1:: Meis1*cKO (Figure 5a, Supplementary Table S1). According to ChIP-Atlas, a data-mining suite for exploring epigenomic landscapes by integrating ChIP-seq (Oki et al., 2018, https://chip-atlas.org), there are 9,599 gene loci where the MEIS1 protein binds in some cells or tissues. Among the 696 genes whose expression is downregulated in *En1:: Meis1*cKO, 355 genes are included in MEIS1-binding genes reported in ChIP-Atlas (Figure 5a, Supplementary Table S2). We refer to these genes as MEIS1 potential target genes, in this study. Interestingly, they include *Vimentin*, *Zeb2*, *Tnc*, and *Slc1a3*, which are known as specific markers of BG-like cells (Figure 5a, Ribotta et al.,2000; He et al.,2018; Bartsch et al., 1992; Tellios et al., 2021). In violin plot data analysis using the scRNA-seq data, it was revealed that the expression of these four genes was reduced in BG-like cells of *En1:: Meis1*cKO cerebella cKO (Figure 5b). Furthermore, immunostaining revealed that expression of VIMENTIN, ZEB2 and TENASCIN (TNC) proteins in astroglial cells (SOX9-, GFAP-double positive cells) was decreased in *En1:: Meis1* cKO cerebella cKO compared to control (Figure 5c, d, e). In the analysis of Figure 5a, *Gria1, Adgrg6* and *Sept4* are not included in the MEIS1-binding gene loci identified by ChIP-Atlas, but it is included in the 696 genes whose expression is downregulated in BG-like cells of *En1:: Meis1*cKO (Figure 5a). Consistently, violin plot showed that its transcript levels were reduced in BG-like cells of the cKO mice (Figure S12a). MEIS1 may increase expression of *Gria1, Adgrg6* and *Sept4* through indirect regulation. These findings suggest that MEIS1 upregulates transcription of BG-like cell specific genes, conferring BG-specific traits to astrocytes (Figure S12b).

**Figure 5.**
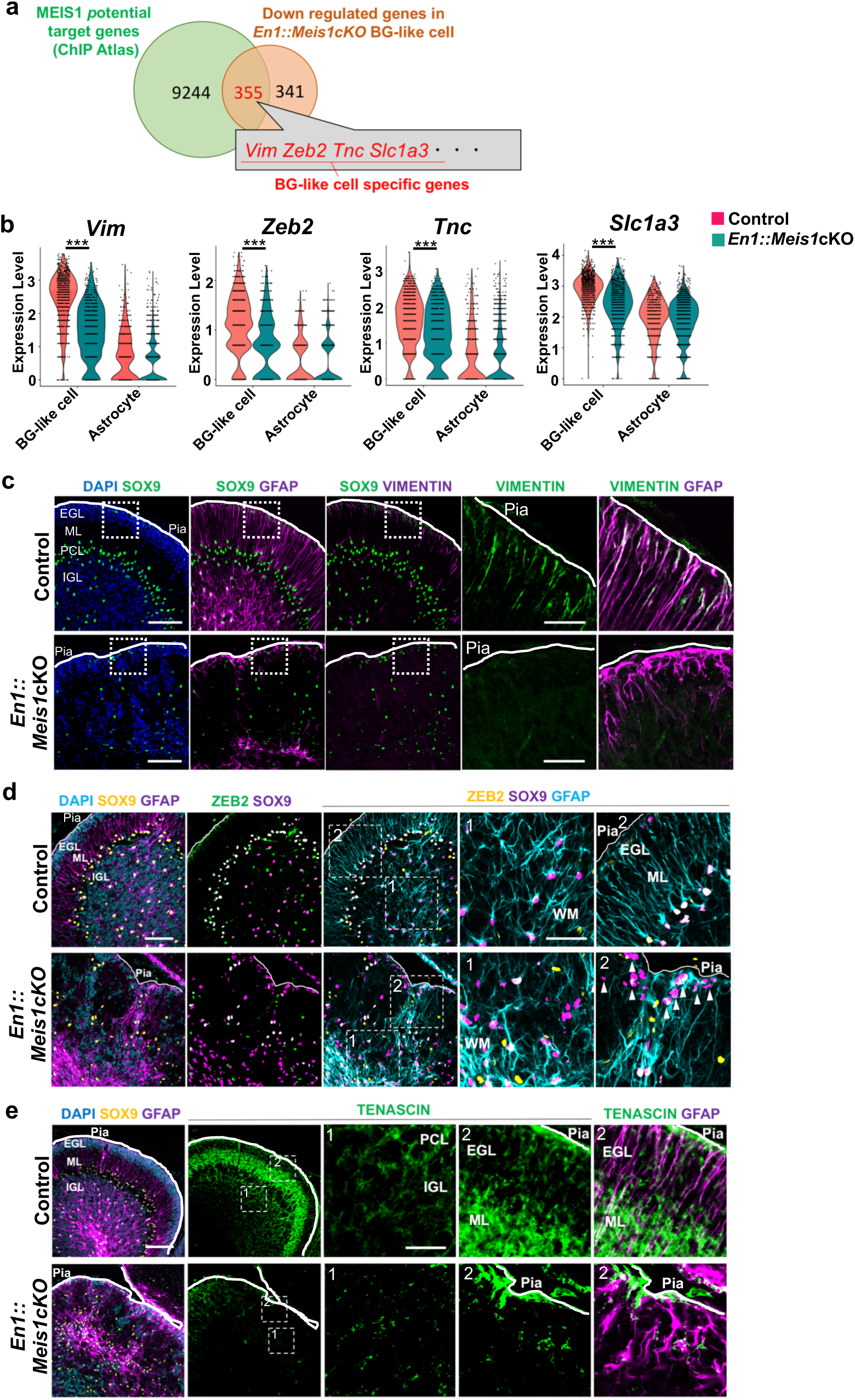
Loss of MEIS1 downregulates the expression of BG-like cell-specific molecules (a) Venn diagram showing the overlap between genes significantly downregulated in BG-like cell subclusters of *En1::Meis1cKO* compared to control (696 genes) and gene loci where the MEIS1 protein binds in some cells or tissues, identified from the ChIP-Atlas database (9,599 genes). 355 genes (MEIS1 potential target genes) were found in common, including BG-like cell–specific genes such as *Vim*, *Zeb2*, *Tnc*, and *Slc1a3*. (b) Violin plots showing the expression levels of BG-like cell–specific genes (*Vim*, *Zeb2*, *Tnc*, and *Slc1a3*) in scRNA-seq data. Expression values were compared between control and *En1::Meis1cKO*, separately for BG-like cells and astrocytes. Statistical analysis was performed using Wilcoxon Rank Sum test. p values are represented as *** for p < 0.001 and * for p < 0.05. (c) Immunostaining of P10 cerebellar sections from control and *En1::Meis1cKO* with SOX9, GFAP, and VIMENTIN. Scale bars: 100 µm (lower magnification), 30 µm (higher magnification). (d) Immunostaining of P10 cerebellar sections from control and *En1::Meis1cKO* with SOX9, GFAP, and ZEB2. Scale bars: 100 µm (lower magnification), 50 µm (higher magnification). (e) Immunostaining of P10 cerebellar sections from control and *En1::Meis1cKO* with SOX9, GFAP, and TENASCIN. Scale bars: 100 µm (lower magnification), 30 µm (higher magnification).

## Discussion

In this study, we demonstrated that MEIS1 is required for BGs, a specialized astrocyte, for the endowment of their characteristic properties in addition to the general features of astrocytes. In *Meis1*-deficient mice, a marked decrease in BGs and increase in multipolar astrocytes were observed, indicating that the distribution of astroglial cells (BGs and astrocytes) became unbalanced. Furthermore, BGLP-specific *Meis1* deletion by *in vivo* electroporation resulted in a pronounced shift in the differentiation direction of BGLP-derived cells from BGs toward astrocytes. In addition, both scRNA-seq and histological analyses suggested that MEIS1 regulates the expression of BG-like cell–specific genes, such as *Zeb2*, *Vim*, *Gria1*, and *Slc1a3*, thereby endowing astrocytes with BG-specific properties. These findings provide the first molecular basis that defines the identity of BGs as a specialized type of astrocyte, and offer a new perspective for understanding astrocyte development.

In both *En1::Meis1cKO* and *Sox9-TX::Meis1cKO* mice, we observed the loss or reduction of unipolar BG-like cells, accompanied by the emergence of ectopically localized astrocytes in the ML and EGL. A similar phenotype characterized by the loss of BG-like cells has been previously reported in mouse models with knockout of *Zeb2*, *Vimentin*, *Ptpn11*, *Huwe1*, and *Sox4*, as well as in mice overexpressing *Mlc1* (He et al., 2018; Ribotta et al.,2000; Li et al., 2014; D’Arca et al., 2010; Hoser et al., 2007; Kikuchihara et al., 2018). Among these genes, *Zeb2* and *Vimentin* were identified in our analysis as BG-like cell (BGLP and BG)-specific genes downstream of MEIS1, suggesting that they may act under MEIS1 to regulate the differentiation from BGLPs to BGs. In addition, *Ptpn11* were included among the *Meis1* potential target genes identified in our *in silico* ChIP-Atlas analysis (Supplementary Table S2). While the expression pattern of *Mlc1* in the cerebellar astroglial lineage had not been fully understood, in our culture experiments of astroglial cells, *Meis1* deletion led to an increase in *Mlc1* expression (Figure 2g). Furthermore, in scRNA-seq analyses, when comparing the gene expression profiles between control and *En1::Meis1cKO* cells in the BG-like cell subcluster, *Mlc1* expression was significantly upregulated in the *En1::Meis1cKO* condition (Supplementary Table S1). Taken together, these findings suggest that the expression of genes previously implicated in the acquisition or maintenance of BG-like cell properties may be directly or indirectly regulated by MEIS1.

We demonstrated that MEIS1 is an essential factor for the differentiation from BGLPs to BGs, and also found that MEIS1 is expressed broadly in astroglial lineage cells, including astrocyte-like cells (AsLPs and astrocytes). Nevertheless, the expression of BG-like cell–specific genes such as *Zeb2*, *Vim*, *Tnc*, and *Slc1a3* was regulated by MEIS1 only in BG-like cells. One possible explanation for this selectivity is that the functional activity of MEIS1 in BG-like cells may require a specific co-factor. Interestingly, in the hematopoietic system, MEIS1 and ZEB2 have been reported to cooperatively activate specific target genes and contribute to cell fate determination (Kitagawa et al., 2023). This raises the possibility that, in the cerebellar astroglial lineage, the expression of BG-like cell–specific genes may depend not on MEIS1 alone, but on its co-expression with ZEB2, one of the BG-like cell–specific genes. This point represents an important issue to be addressed in future studies.

In *En1::Meis1cKO* mice, disruption of BG-like cell processes was already observed at P3, preceding the disorganization of normal cortical layers and the mispositioning of neurons. During cerebellar development, it has been reported that the loss of BG-like cell processes markedly impairs GC migration and layer formation during the first two postnatal weeks, leading to ataxia (Cerrato et al., 2020). Considering that the structural abnormalities and reduction of cerebellar volume became more pronounced within the first postnatal week in *En1::Meis1cKO* mice, it is conceivable that at least part of the phenotypes of *En1::Meis1cKO*, including impaired motor learning, are induced by the early loss of BG-like cells caused by the failure of differentiation from postnatal BGLPs into BGs.

Interestingly, the loss of BG-like cells and other phenotypes observed in *En1::Meis1cKO* mice were more prominent in the rostral cerebellum, whereas they appeared relatively mild in the caudal region. A similar rostrocaudal gradient of phenotypes has also been reported in the cerebellum of mice in which *Zeb2* was deleted in astroglial lineage cells (He et al., 2018). These observations suggest that the molecular mechanism underlying the endowment of BG-specific properties, regulated by *Meis1* and its downstream target *Zeb2*, may be preferentially activated in the rostral region of the cerebellum. Further studies will be required to elucidate the regulatory mechanisms that control BG differentiation in the caudal cerebellar region.

In this study, we presented the molecular basis by which astrocytes are endowed with the characteristic properties of BGs and identified a key factor essential for the differentiation from BGLPs to BGs. Furthermore, our findings suggest that the reduction or loss of BGs caused by MEIS1 deficiency may lead to cerebellar malformation and ataxia. Taken together, the molecular mechanisms controlling cerebellar astrocyte development centered on MEIS1 will be of great significance not only for developmental but also for clinical research.

## Experimental procedures

### Animals

All animal experiments were approved by the Animal Care and Use Committee of the National Institute of Neuroscience, National Center of Neurology and Psychiatry (Tokyo, Japan; Project 202017R2). The *Atoh1-Cre-T*g, *En1^Cre^*, *Sox9^CreERT2/+^* and *Meis1 ^fl^* mouse lines have been described previously (Fujiyama et al., 2009; Kimmel et al., 2000; Soeda et al., 2010; Ariki et al., 2014). Wild-type and genetically modified mice were maintained on a C57BL/6 background. For tamoxifen and EdU administration, *Meis1^fl/fl^,* and *Meis1^fl/fl^; Sox9^CreERT2/+^*mice received intraperitoneal injections of tamoxifen (10 mg/kg, SIGMA) and EdU (5 mg/kg, Thermo Fisher Scientific) at P0 and P1. Mice were housed under specific pathogen-free (SPF) conditions with a 12-h light/dark cycle and allowed to freely access to food and water. Both male and female mice were used in this study.

### Perfusion and tissue preparation

Mice younger than P10 were euthanized by decapitation, and brains were immediately dissected and fixed overnight in 4% paraformaldehyde (PFA) in PBS at 4 °C with gentle agitation. Mice older than P10 were transcardially perfused with PBS followed by 4% PFA, after which the brains were removed and post-fixed in 4% PFA overnight at 4 °C. Fixed brains were cryoprotected sequentially in 1× PBS, 10% sucrose, and 30% sucrose (each overnight at 4 °C with gentle shaking). Cryoprotected tissues were embedded in O.C.T. compound (Sakura Finetek), rapidly frozen, and sagittally sectioned at 16 µm thickness using a cryostat (CM3050S; Leica). For the bright-field macroscopic observation shown in Figure 1a, P21 brains were perfusion-fixed, removed from the skull, and imaged prior to cryoprotection and sectioning.

### Immunohistochemistry and antibodies

Cryosections were washed twice with PBS for 10 min and then incubated at room temperature with 10% normal donkey serum (S30-100ML; Millipore) containing 0.2% PBST (Triton) for 40 min. After blocking, the sections were incubated with primary antibodies diluted with blocking solutions at 4 °C for 16 h. The following primary antibodies were used: mouse anti-Calbindin D28K (1:500; Sigma-Aldrich), rat anti-GFAP (1:500; Merck Millipore), chicken anti-GFP (1:1000; Abcam), rat anti-KI67 (1:500; Invitrogen), goat anti-NeuroD1 (1:500; R&D Systems), goat anti-SOX9 (1:500; R&D Systems), rabbit anti-Vimentin (1:500; Cell Signaling Technology), rabbit anti-MEIS1 (1:500; homemade; Owa et al., 2018), rabbit anti–cleaved Caspase-3 (1:100; Cell Signaling Technology), rabbit anti-ZEB2 (1:500; Sigma-Aldrich) and rabbit anti-TENASCIN (1:500; Merck Millipore). After primary antibody incubation, sections were washed twice with PBS (10 min each) and incubated for 2 h at room temperature with species-appropriate secondary antibodies conjugated to Alexa Fluor 405, 488, 568, or 647 (1:400; Abcam or Jackson ImmunoResearch), together with DAPI (25 µg/mL; Invitrogen) in 0.2% PBST. Following two additional PBS washes (10 min each), samples were mounted with ProLong Glass Antifade Mountant (P36984; LTJ). For EdU detection, sections were processed using the Click-iT™ EdU Alexa Fluor™ 647 Flow Cytometry Assay Kit (Thermo Fisher Scientific) after the secondary antibody incubation and subsequent PBS washes, according to the manufacturer’s instructions (used here for tissue sections).

### Nissl staining

Frozen sections were air-dried and immersed for 7 min in an organic solvent mixture of chloroform: diethyl ether: ethanol (1 : 1 : 7). The sections were then rehydrated sequentially in 100%, 70%, and 50% ethanol, followed by staining in 0.1% cresyl violet solution containing 1% acetic acid. After staining was visually confirmed, the sections were dehydrated through 50%, 70%, and 100% ethanol, cleared in xylene, and coverslipped using a mounting medium (Merck Millipore).

### Primary cerebellar astroglial culture

Primary astroglial cultures were prepared from neonatal mouse cerebella following previously described procedures with minor modifications (Tellios et al., 2021; Martínez-Lozada et al., 2011). Briefly, 200 µL of Leibowitz L-15 medium containing 100 mg/mL bovine serum albumin (BSA) and 1× penicillin/streptomycin (P/S) was placed on ice in a 1.5 mL microcentrifuge tube. Neonatal mice (P0–P2) were euthanized by decapitation, and cerebella were rapidly dissected and transferred into the chilled L-15 medium. Tissue was mechanically dissociated first with a 1 mL pipette and subsequently with a 200 µL pipette to obtain a homogenate, which was then passed through a 70 µm nylon cell strainer. The flow-through was centrifuged, and the resulting pellet was resuspended in DMEM supplemented with 10% fetal bovine serum (FBS) and 1× P/S. Cells derived from one cerebellum were plated onto one well of a 6-well plate (growth area 9 cm²). Cultures were maintained for 7–10 days with medium changes every 3 days, until reaching confluence. Confluent astroglial lineage cells were then trypsinized and replated at a density of 5×10^4^ cells per well (growth area 9 cm²) for downstream assays. For immunocytochemistry (ICC), sterile coverslips (3–4 per well) were placed in the wells prior to replating. Cells were fixed for ICC or harvested for RNA extraction at either 3 or 7 day after replating. For experiments requiring *Meis1* deletion, 4-hydroxytamoxifen (4-OH-TX) was added to cultures at 5 or 10 µM (vehicle: methanol) 3 days after replating, and cells were fixed or collected 4 days later.

### Immunocytochemistry and antibodies

Cells cultured on coverslips were fixed in 4% paraformaldehyde (PFA) for 10–15 min at room temperature, rinsed with 1× PBS, and stored at 4 °C until further processing. Fixed cells were washed twice with 1 mL of PBS (5 min each) and permeabilized with 1 mL of 0.1% PBST (PBS containing 0.1% Triton X-100) for 10 min at room temperature. After removal of the permeabilization buffer, cells were incubated in blocking solution consisting of 10% normal donkey serum (NDS) in 0.1% PBST for 30 min at room temperature. Primary antibodies were diluted in blocking solution and applied to the cells (500 µL per well), followed by incubation at 4 °C overnight. The following primary antibodies were used: rat anti-GFAP (1:1000; Merck Millipore), rabbit anti-GLAST (1:1000; Abcam), rat anti-KI67 (1:1000; Invitrogen), and rabbit anti-MEIS1 (1:1000; homemade; Owa et al., 2018). After primary antibody incubation, cells were washed three times with 1 mL of PBS (5 min each). Secondary antibodies conjugated to Alexa Fluor dyes and DAPI were diluted in 0.1% PBST and applied to the cells (500 µL per well; secondary antibodies 1:1000; Abcam or Jackson ImmunoResearch, DAPI 1:2000; Invitrogen) for 1 h at room temperature. Following three additional PBS washes (5 min each) and a brief rinse in Milli-Q water, coverslips were mounted using ProLong Glass Antifade Mountant (LTJ).

### cDNA synthesis and quantitative PCR (qPCR)

Total RNA was extracted from primary cerebellar astroglial cultures using the RNeasy Plus Micro Kit (Qiagen) according to the manufacturer’s instructions. Reverse transcription was performed using the ReverTra Ace qPCR RT Kit (Toyobo). RNA samples were heated at 65 °C for 5 min and immediately placed on ice. Each 10 µL RT reaction contained 2 µL of 5× RT Buffer, 0.5 µL RT Enzyme Mix, 0.5 µL Primer Mix, RNA template, and nuclease-free water to volume. The reverse transcription reaction was carried out at 37 C for 15 min, followed by enzyme inactivation at 98 °C for 5 min. Both RT-positive (RT+) and RT-negative (RT–) reactions were prepared and stored at 20 °C until use. qPCR reactions (10 µL total volume) consisted of 5 µL SYBR Green Master Mix (Thermo Fisher Scientific), 2 µL primer mix (0.5 µM each of forward and reverse primers), 1 µL cDNA, and 2 µL nuclease-free water. All reactions were run in technical triplicate on a LightCycler 96 system (Roche). For each gene, relative expression levels were calculated using the vehicle-treated (MeOH) samples as the reference, and fold changes in the 4-hydroxytamoxifen (4-OH-TX)–treated samples were quantified using the ΔΔCt method. Primer sequences for all genes, including the housekeeping gene *Gapdh*, are provided in Supplementary Table S3.

### Image acquisition and quantification

Nissl-stained sections were imaged using an all-in-one fluorescence microscope (BZ-X700; Keyence). Confocal images of brain sections (IHC) and cultured cells (ICC) were acquired using a spinning-disk confocal microscope (SpinSR10; Olympus). For IHC analyses, images were collected from rostral lobules I–III and caudal lobules VIII–X of the cerebellum. For experiments involving *in vivo* electroporation, lobules IV–VI of the cerebellar vermis were examined. For each analysis, at least two non-overlapping images were acquired per animal. All acquired images were processed and quantified using ImageJ (NIH), and graphs were generated using GraphPad Prism (GraphPad Software).

#### In vivo electroporation

*In vivo* electroporation in neonatal mice was performed as previously described (Suyama et al., 2025, bioRxiv; Miyashita et al., 2021; Adachi et al., 2021; Yamashita et al., 2020; Owa et al., 2018). Expression plasmids were diluted to 1 µg/µL in Milli-Q water, and 0.02% Fast Green was added to visualize the solution. P0 pups were anesthetized on ice, and 10 µL of plasmid solution was applied onto the surface of the cerebellar vermis. Electroporation was performed using a forceps-type electrode (CUY650P5-3, NEPA GENE) with 70 V square pulses (pulse duration: 50 ms; pulse interval: 150 ms; seven pulses). Following electroporation, pups were maintained on a 37 °C warming plate until fully recovered and then returned to their litter. This procedure consistently introduced genes into lobules IV–VI of the cerebellar vermis.

#### Rotarod test

The rotarod test was performed as previously described (Yamashiro et al., 2020). Briefly, motor coordination and motor learning were assessed using an accelerating rotarod apparatus (Rotarod 47600; Ugo Basile). Two-month-old mice were placed on the rod at 4 rpm for 10 s, after which the rotation speed was linearly accelerated from 4 to 40 rpm over 300 s. Mice underwent three trials per day for two consecutive days, with at least 10-min intervals between trials. For each trial, the latency to fall—or to cling to the rotating rod and complete a full passive rotation—was recorded.

#### Single-cell RNA Sequencing

Single-cell RNA sequencing was performed on eight samples in total, comprising two genotypes (*Meis1^fl/fl^* and *Meis1^fl/fl^; En1^Cre/+^*) and two developmental stages (P3 and P10), with n = 2 biological replicates per condition. Single-cell RNA-seq libraries were prepared by MACROGEN and GENEWIZ (Azenta Life Sciences). Libraries were sequenced on an Illumina platform (HiSeq, instrument ID: E00516A; paired-end mode) according to the provider’s standard protocols. Sequence files obtained from the second batch were split into two independent sequencing datasets per sample. Consequently, a total of 12 sequencing datasets were generated (n = 3 datasets per condition, N = 2 biological replicates per condition) and used for downstream analyses. Data analysis was conducted in R (v4.4.2). Normalization, dimensionality reduction, clustering, and visualization of scRNA-seq data were performed by Seurat (v5.3.0, Satija et al, 2015) function of NormalizeData, RunPCA, RunUMAP, FindNeighbors, FindCluster, FindAllMarkers with default parameters.

#### In silico ChIP-Atlas Analysis

*In silico* identification of putative MEIS1 target genes was performed using ChIP-Atlas (https://chip-atlas.org; Oki et al., 2018). The “Target Genes” function was applied with the Mus musculus (mm10) dataset, using MEIS1 as the query factor and a search window of ±10 kb from the transcription start site (TSS). Only genes with an average MEIS1 score exceeding 20 were retained for further analysis. This analysis yielded 9,599 genes that were potentially regulated by MEIS1.

#### Statistical Analysis

Individual animals or trials were treated as biological replicates, and all mice were handled under identical experimental conditions. Data are presented as mean ± SEM. Statistical analyses were performed using GraphPad Prism (GraphPad Software). Student’s t-tests, Tukey’s multiple comparisons test, Dunnett’s multiple comparisons test and Wilcoxon Rank Sum test were applied as appropriate. Statistical significance is indicated as follows: n.s., p > 0.05; *p < 0.05; **p < 0.01; ***p < 0.001; ****p < 0.0001.

## Acknowledgements

This work was supported by JSPS KAKENHI (Grant Numbers JP22H02730 to MH, 22K15211 to T.O, 21K20853 and 23K14203 to TA); AMED (Grant Numbers 24wm0425005h0004 and 24ek0109764h0001 to MH), an Intramural Research Grant of NCNP (4–5, 4-6, 6-9 to MH); Japan Health Research Promotion Bureau (JH) under Research Fund (2020-B-07 and 2024-D-01 to MH); Multilayered Stress Diseases (JPMXP1323015483 to MH); Tokumori Yasumoto Memorial Trust (MH) and Takeda Science Foundation (Grants 2024049458 to TA).

## Author contributions

Conceptualization: T.A., T.O., and M.H. Methodology: K.I., T.A., T.O., and M.H. Investigation: K.I., T.A., T.O., M.M., K.S., K.J., K.H., I.H., E.I., and S.M. Resources: Y.U.I., R.G., T.N., and T.I. Formal analysis: K.I., and T.A. Writing—original draft: T.A. and M.H. Writing—review and editing: All authors. Visualization: K.I., T.A., and K.S. Funding acquisition: T.A., T.O., and M.H. Supervision: T.A., T.O., M.S., K.K., T.Y., and M.H.

## Data availability

The transcriptomics data is deposit in the Zenodo (10.5281/zenodo.18308841) and is publicly available as of the date of publication.

## Code availability

The custom code associated with this study is available at https://github.com/Hoshino-lab/Ichijo_2026_scRNAseq.

---

**Figure S1.**
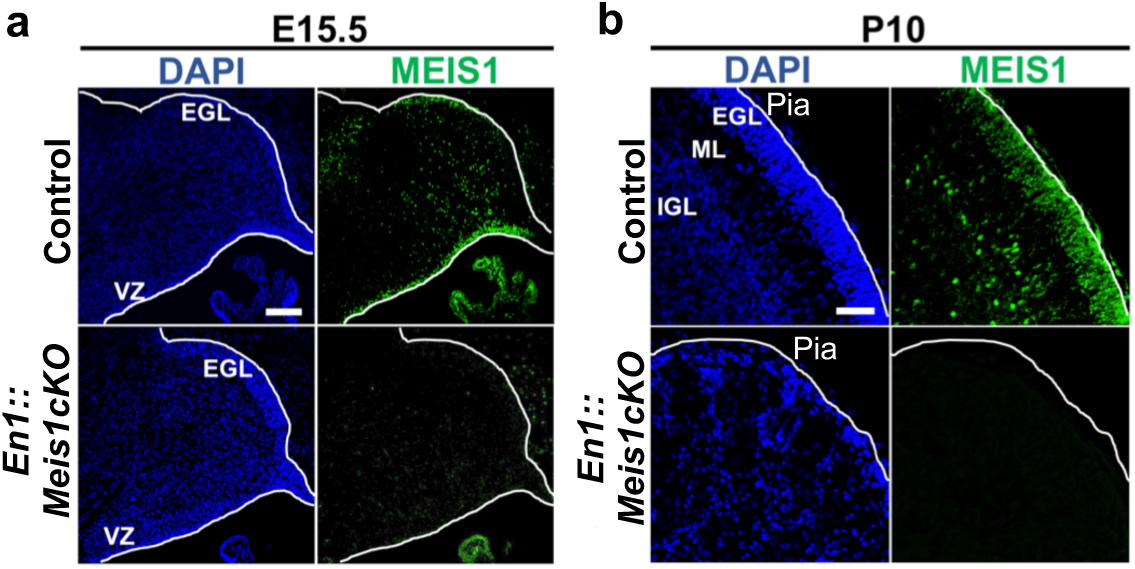
MEIS1 protein is lost in the cerebellum of *En1::Meis1cKO* (a, b) Immunostaining of cerebellar sections from control and *En1::Meis1cKO* with MEIS1 antibody at E15.5 (a) and P10 (b). Scale bars: 50 µm (a), 100 µm (b).

**Figure S2.**
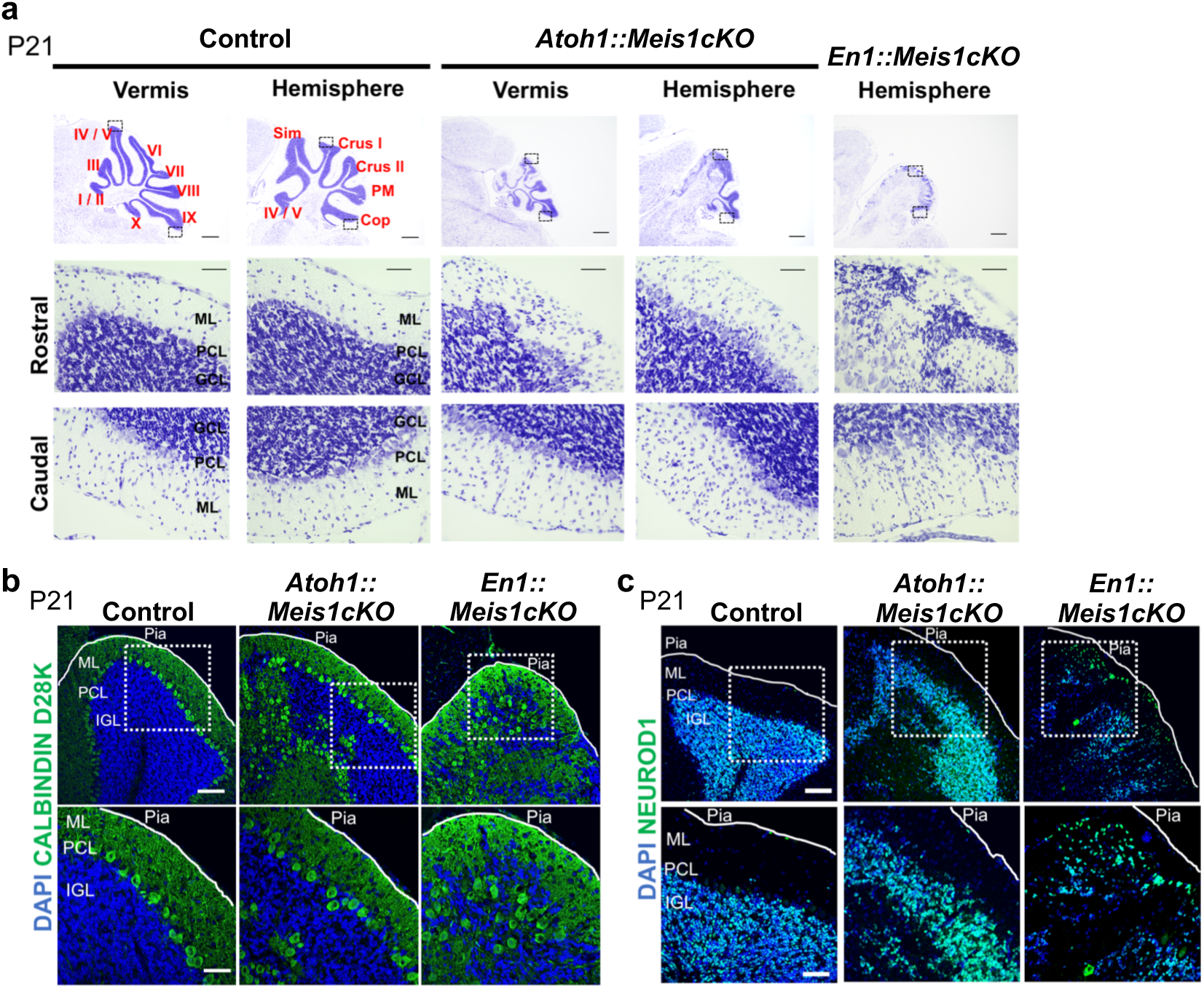
Loss of *Meis1* in the whole cerebellum leads to cerebellar layer disorganization (a) Nissl-stained cerebellar sections from control, *Atoh1::Meis1cKO*, and *En1::Meis1cKO* at P21. Both vermis and hemisphere are shown for control and *Atoh1::Meis1cKO*, whereas only the hemisphere is displayed for *En1::Meis1cKO* due to the absence of vermis region. Higher-magnification images of rostral and caudal regions are also shown in each genotype. Scale bars: 500 µm for low-magnification images, and 50 µm for high-magnification images. (b, c) Immunostaining of P21 cerebellar sections from control, *Atoh1::Meis1cKO*, and *En1::Meis1cKO* with (b) CALBINDIN-D28k (Purkinje cells) and (c) NEUROD1 (granule cells). Scale bars: 50 µm for lower magnification and 25 µm for higher magnification.

**Figure S3.**
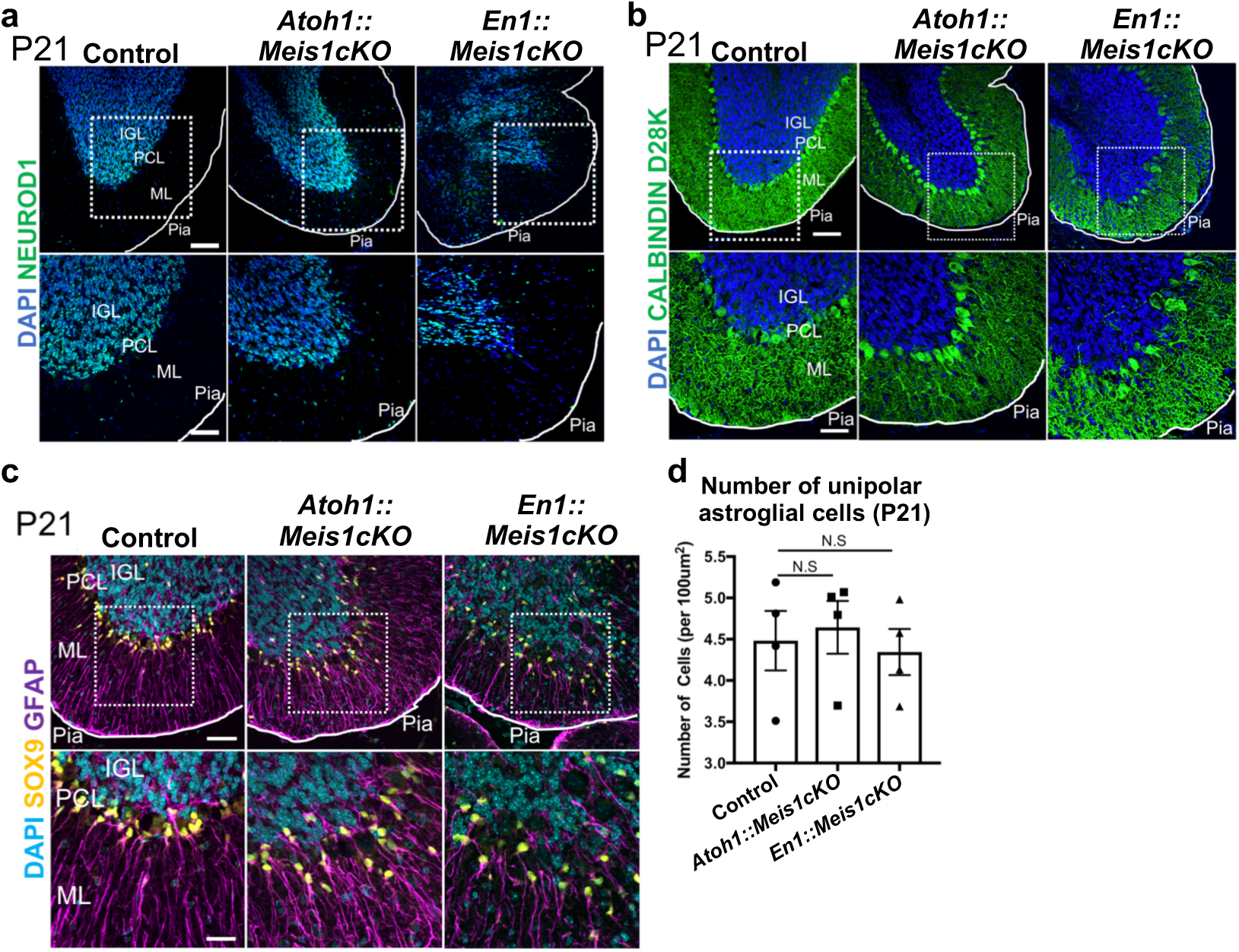
The phenotype of *Meis1* cKO mice is milder in the caudal cerebellum (a–c) Immunostaining of P21 cerebellar sections from the caudal region of control, *Atoh1::Meis1cKO*, and *En1::Meis1cKO* with (a) NEUROD1 (granule cells), (b) CALBINDIND28k (Purkinje cells), and (c) SOX9 and GFAP (astroglial cells). Scale bars: 50 µm for upper panels and 25 µm for lower panels. (d) Quantification of unipolar astroglial cells (BG-like cells) in the caudal cerebellum at P21. Statistical analysis was performed using Dunnett’s multiple comparisons test. N.S. indicates p > 0.05.

**Figure S4.**
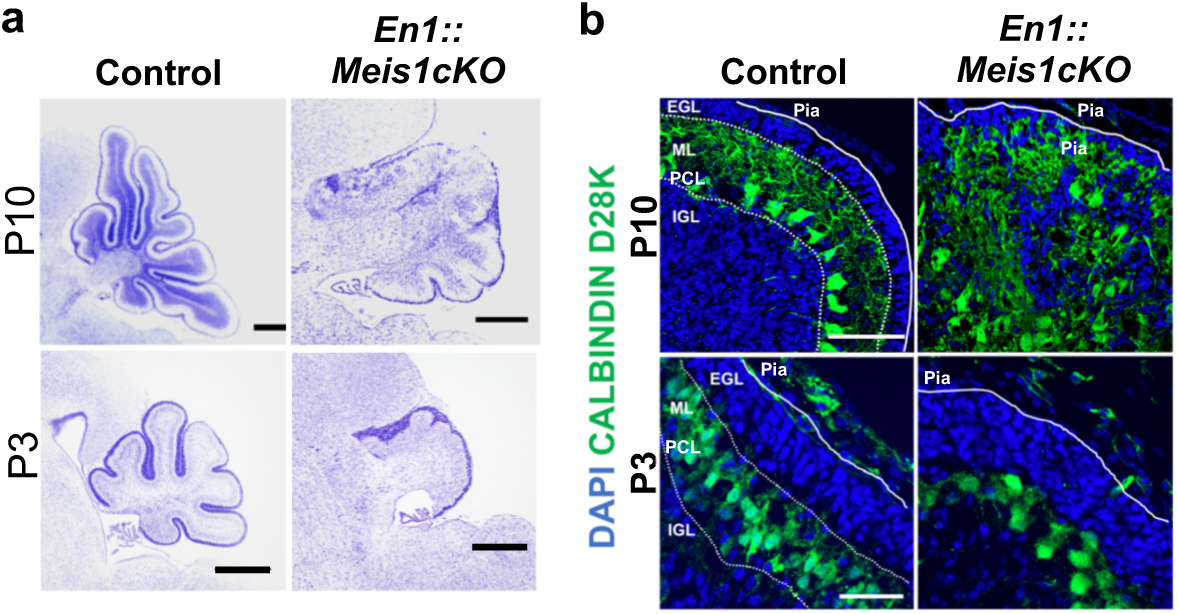
Impaired postnatal cerebellar growth and Purkinje cell mislocalization in *En1::Meis1cKO* (a) Nissl staining of cerebellar sections from control and *En1::Meis1cKO* at P3 and P10. Scale bars: 300 µm (P10 control), 200 µm (P10 *En1::Meis1cKO*, P3). (b) Immunostaining with CALBINDIND28k (Purkinje cells) in control and *En1::Meis1cKO* at P3 and P10. At P10, Purkinje cells were ectopically localized, whereas at P3 their positioning was relatively preserved. Scale bars: 100 µm (P10), 50 µm (P3).

**Figure S5.**
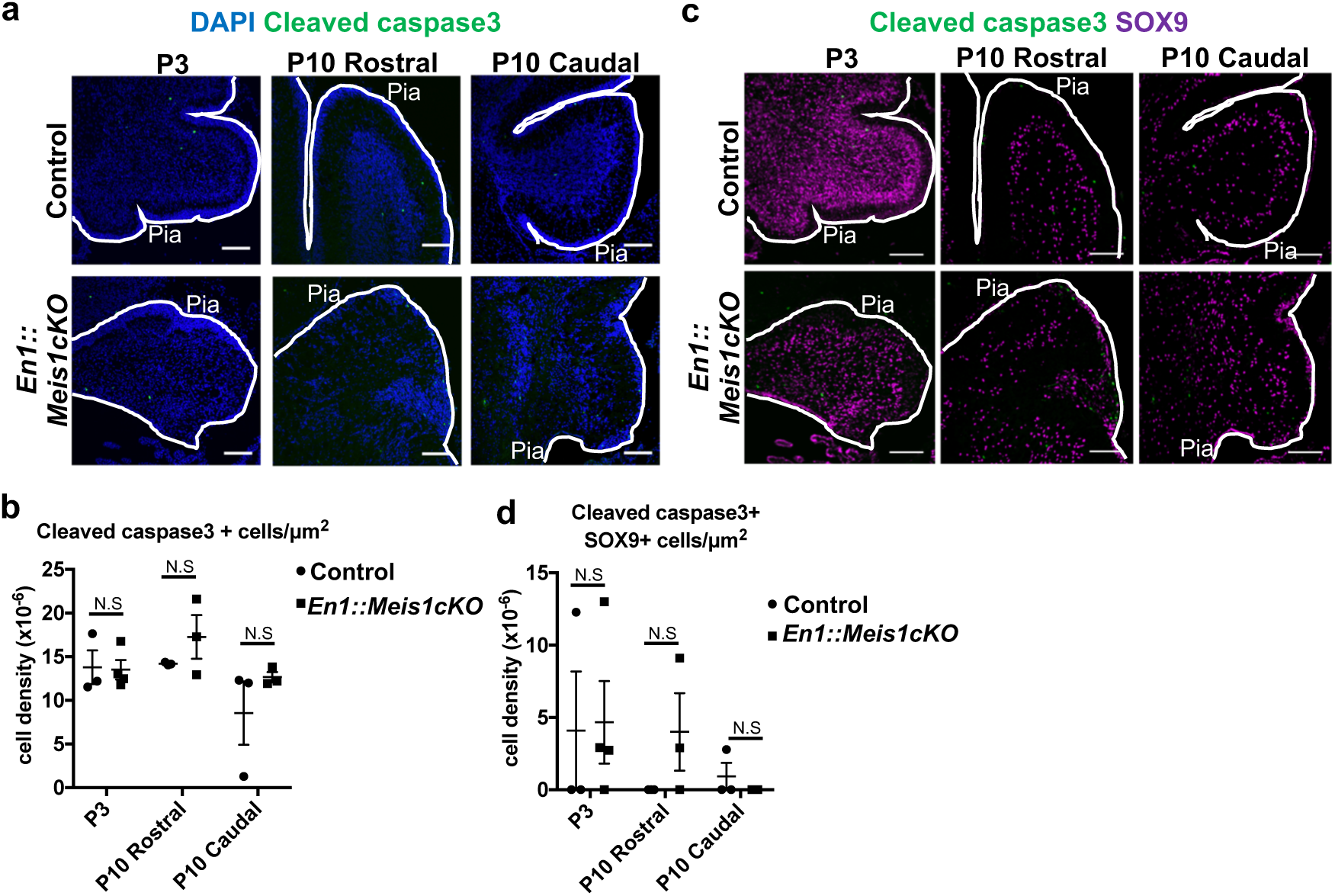
The loss of *Meis1* does not increase apoptosis during the postnatal stage (a) Immunostaining of cleaved caspase-3 (apoptotic cell marker) in the cerebella of control and *En1::Meis1cKO* at P3 and P10. Scale bars: 100 µm. (b) Quantification of cleaved caspase-3–positive cell density at P3 and P10 in control and *En1::Meis1cKO* cerebella. Statistical analysis was performed using unpaired t-tests; N.S. indicates p > 0.05. (c) Immunostaining of cleaved caspase-3 and SOX9 (astroglial cell marker) in the cerebella of control and *En1::Meis1cKO* at P3 and P10. Scale bars: 100 µm. (d) Quantification of cleaved caspase-3+ and SOX9+ cell density at P3 and P10 in control and *En1::Meis1cKO* cerebella. Statistical analysis was performed using unpaired t-tests; N.S. indicates p > 0.05.

**Figure S6.**
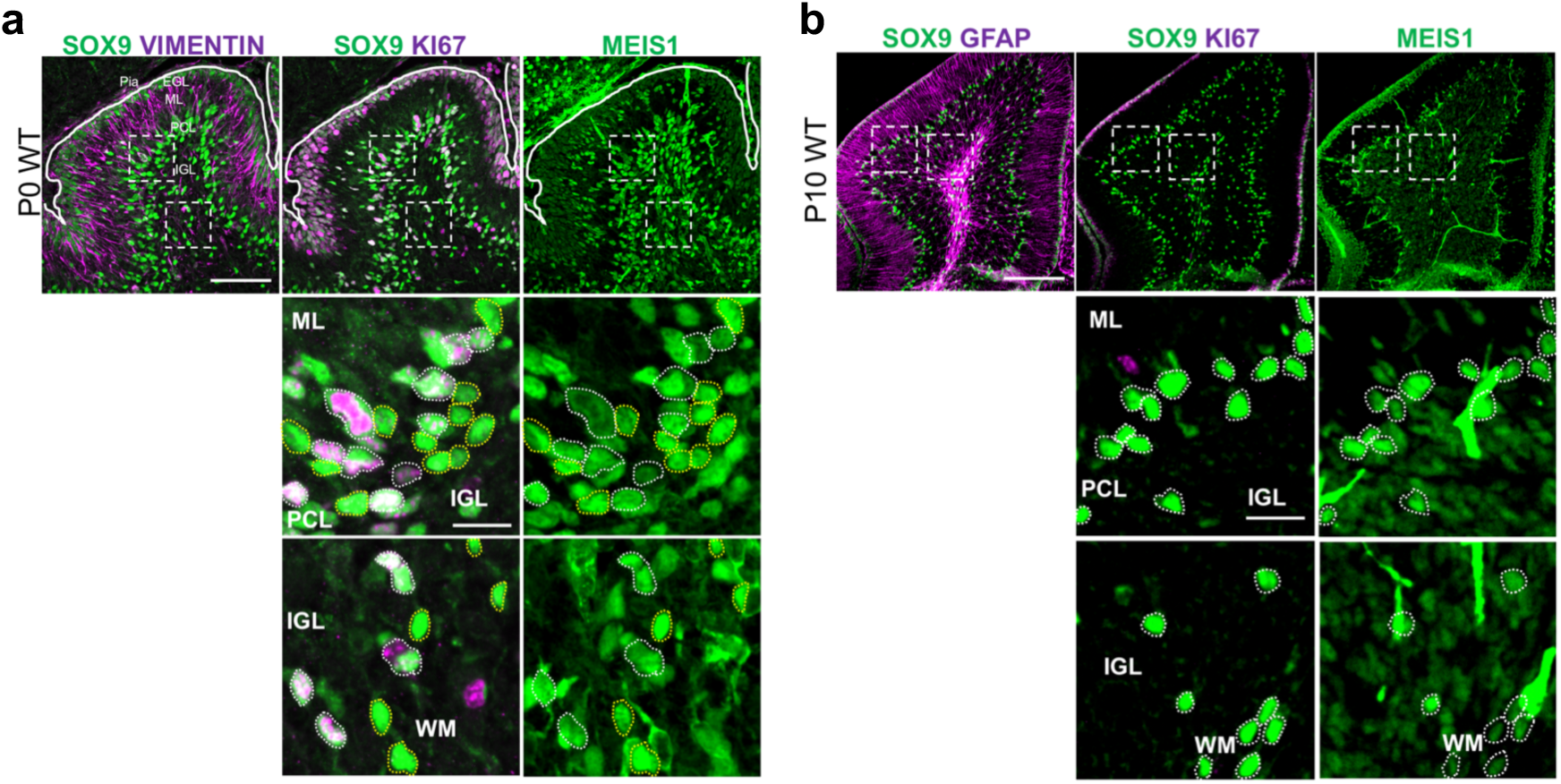
Expression profiles of MEIS1 in developing cerebellar astroglial lineage cells (a, b) Immunostaining of cerebellar sections at (a) P0 and (b) P10 with MEIS1 and astroglial lineage markers (SOX9 and VIMENTIN at P0; SOX9 and GFAP at P10), together with the proliferative marker KI67. Scale bars: 100 µm (lower magnification), 20 µm (higher magnification).

**Figure S7.**
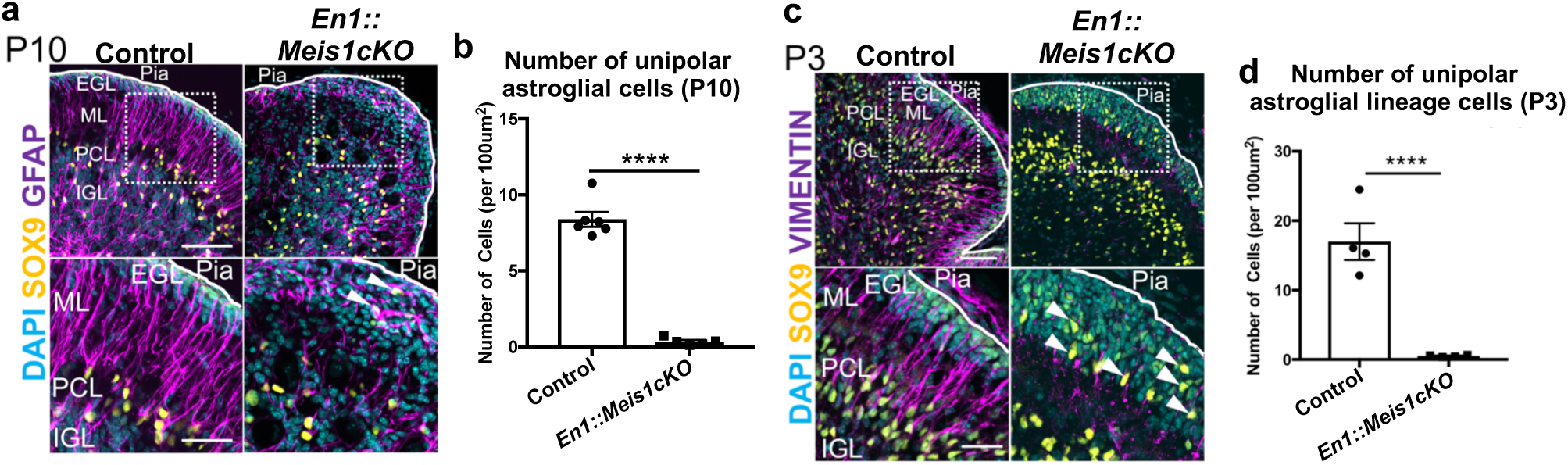
Loss of BG-like cells in the developing postnatal cerebellum of En1::Meis1cKO (a) Immunostaining for SOX9 and GFAP (astroglial cells) in control and *En1::Meis1cKO* cerebella at P10. White arrowheads indicate ectopically localized multipolar astrocytes near the cerebellar surface. Scale bars: 100 µm (upper panels), 50 µm (lower panels). (b) Quantification of the unipolar astroglial cell (BG-like cell) number in the P10 cerebellum of each genotype. Statistical analysis was performed using an unpaired t-test. **** indicates p < 0.0001. (c) Immunostaining for SOX9 and VIMENTIN (astroglial cells) in control and *En1::Meis1cKO* cerebella at P3. White arrowheads indicate ectopically localized multipolar astrocytes near the cerebellar surface. Scale bars: 100 µm (upper panels), 50 µm (lower panels). (d) Quantification of the unipolar astroglial-lineage cell (BG-like cell) number in the P3 cerebellum of each genotype. Statistical analysis was performed using an unpaired t-test. **** indicates p < 0.0001.

**Figure S8.**
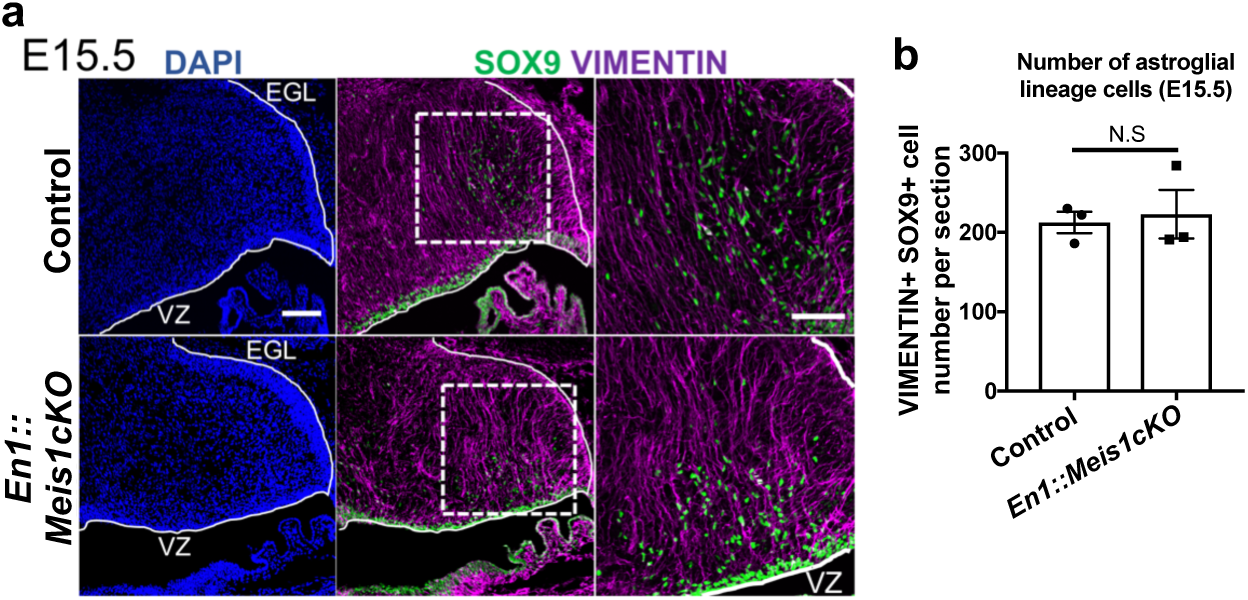
The loss of *Meis1* does not alter the number of astroglial lineage cells during the embryonic stage(a) Immunostaining of the cerebellum at E15.5 from control and *En1::Meis1cKO* with SOX9 and VIMENTIN (astroglial lineage cells). Scale bars: 100 µm (lower magnification), 50 µm (higher magnification). (b) Quantification of SOX9 and VIMENTIN double-positive cells (astroglial lineage cells) that had migrated away from the ventricular zone (VZ). Statistical analysis was performed using an unpaired t-test. N.S. indicates p > 0.05.

**Figure S9.**
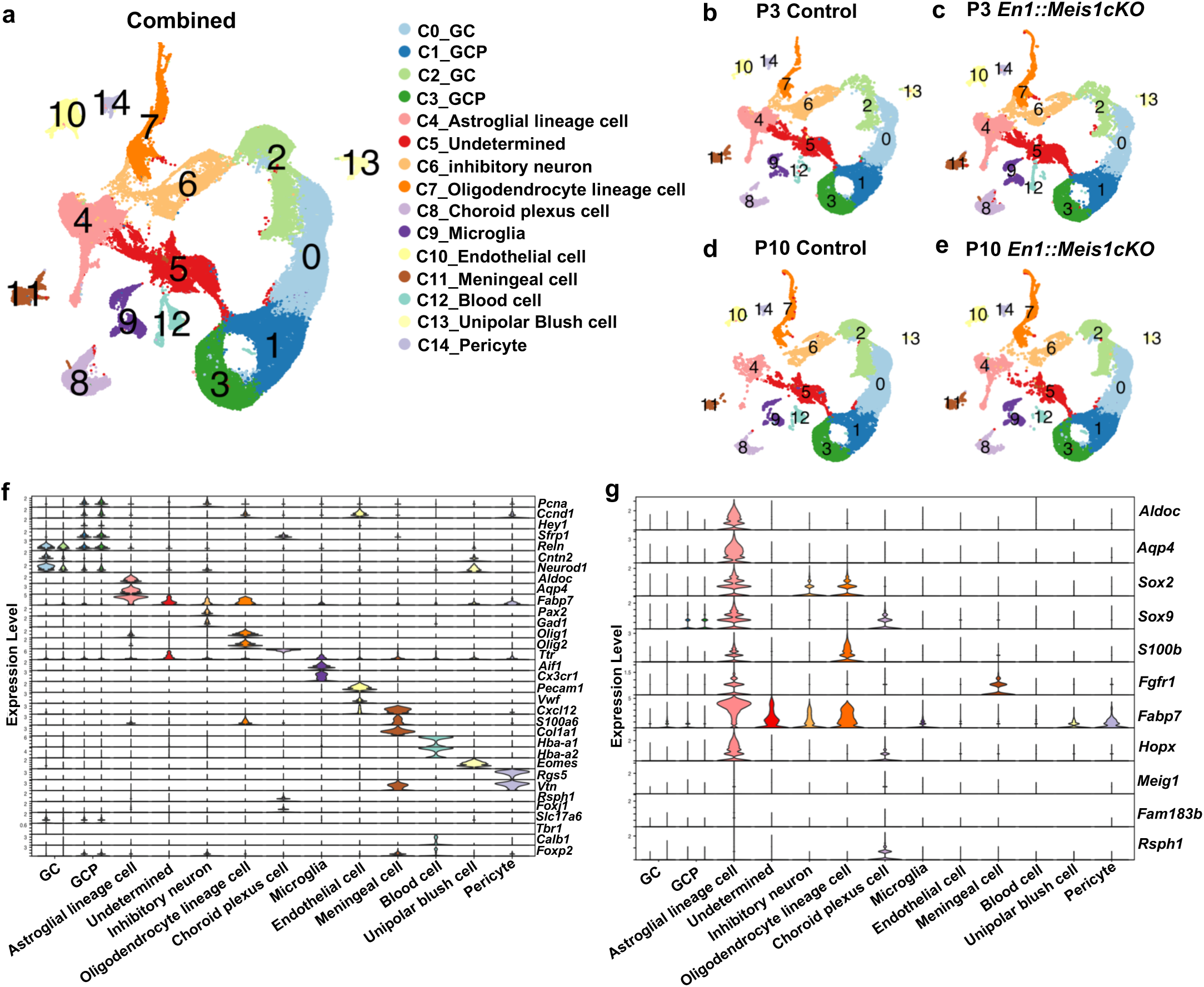
Supplemental data for single cell RNA-seq (a–e) UMAP dimensional reduction of all cells (79,038 cells). Plots show all four condition combined (a), P3 control (b), P3 *En1::Meis1cKO* (c), P10 control (d), and P10 *En1::Meis1cKO* (e). (f) Violin plots showing the expression of known marker genes corresponding to distinct cerebellar cell types across the 15 identified clusters. (g) Violin plots showing the expression of common astroglial lineage markers (*Aldoc, Aqp4, Sox2, Sox9, S100b, Fgfr1, Fabp7, Hopx*) and ciliated cell (Essen et al., 2020) markers (*Meig1, Fam183b, Rsph1*) across the 15 identified clusters.

**Figure S10.**
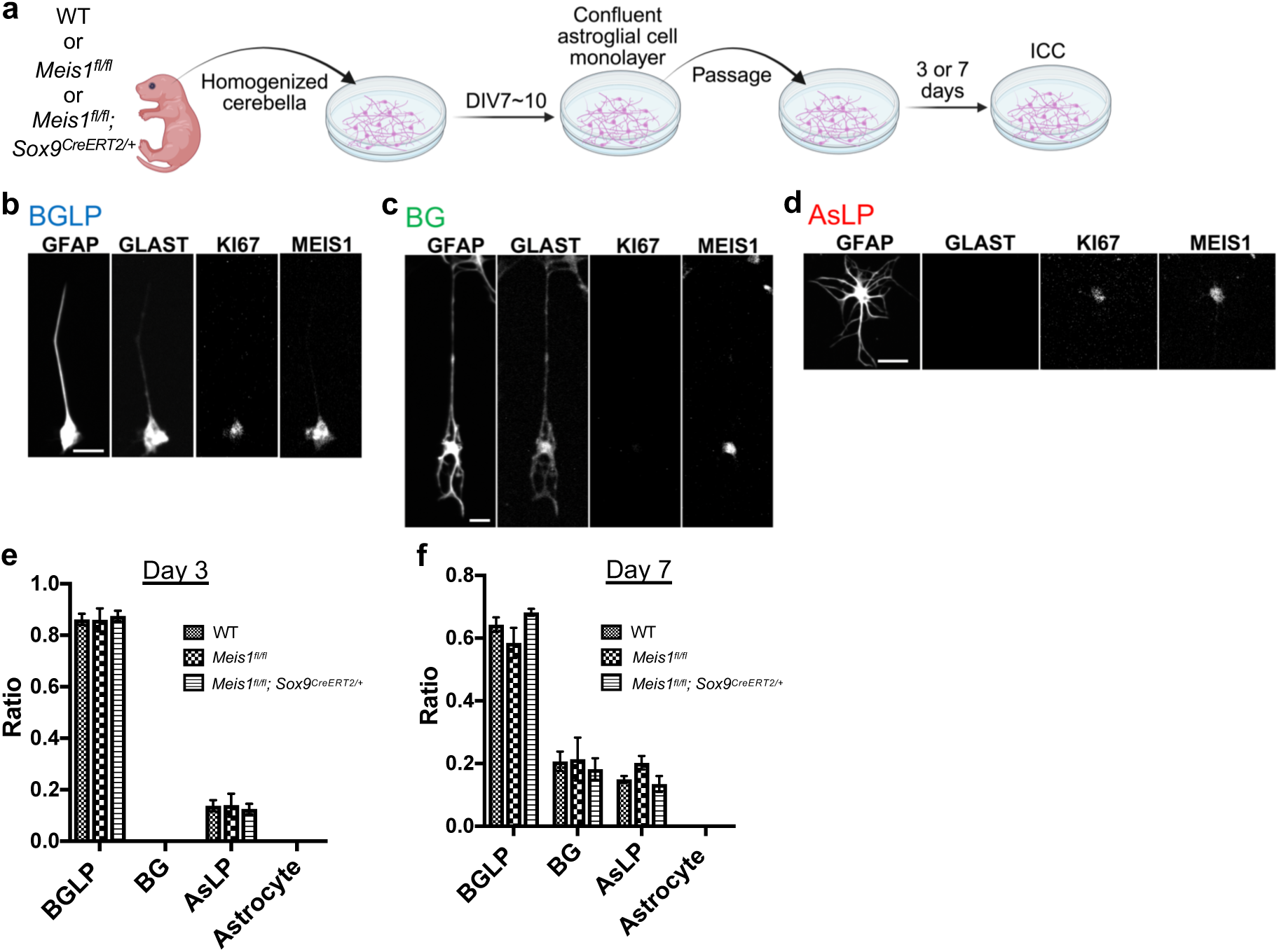
Supplemental data for Figure 2 (a) Experimental scheme. Cerebella were isolated from WT, *Meis1^fl/fl^*, and *Meis1^fl/fl^; Sox9^CreERT2/+^* mice, dissociated, and cultured. When cultures reached confluence at DIV7–10, cells were passaged at a density of 1×10^5^ cells per well. Immunocytochemistry (ICC) was performed either 3 days or 7 days after passage. (b–d) Immunocytochemistry of cultured cerebellar astroglial cells with GFAP, GLAST, KI67, and MEIS1. BG-like progenitors (BGLPs), characterized by GLAST and KI67 double positivity and a unipolar morphology (b). Bergmann glial cells (BGs), characterized by GLAST positivity, KI67 negativity, and a unipolar morphology (c). And Astrocyte-like progenitors (AsLPs), characterized by the absence of GLAST expression, KI67 positivity, and a multipolar morphology (d). Differentiated astrocytes (KI67 negative, GLAST negative and multipolar morphology) were not observed under these conditions. All astroglial lineage cells examined consistently expressed MEIS1. Scale bar, 10 µm. (e, f) Quantification of astroglial lineage cell ratios in cultures derived from the three genotypes at different fixation time points. Cultures were fixed at day 3 (e) or day 7 (f) after passage. Statistical analysis was performed using Dunnett’s multiple comparisons test, and no significant differences were detected among genotypes (p > 0.05).

**Figure S11.**
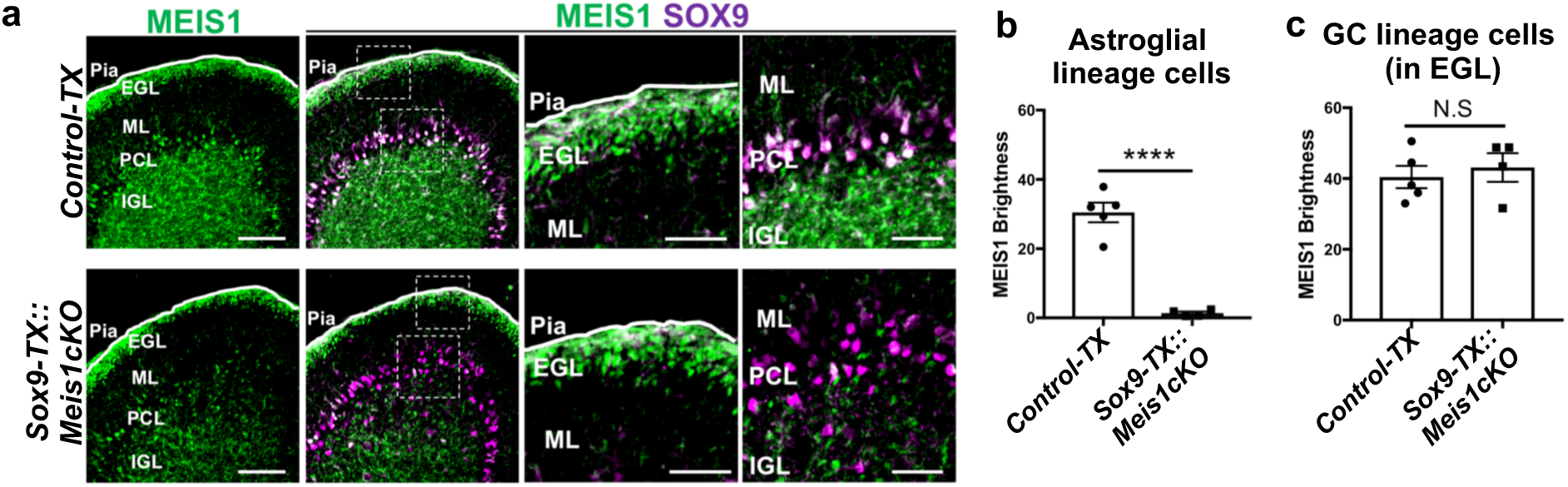
Supplemental data for Figure 3 (a) Immunostaining of P10 cerebellar sections from *Control-TX* and *Sox9-TX::Meis1cKO* with MEIS1 and SOX9. Scale bars: 50 µm (lower magnification), 25 µm (higher magnification). (b, c) Quantification of MEIS1 fluorescence intensity in astroglial lineage cells (b) and in GC lineage cells within the EGL (c) from *Control-TX* and *Sox9-TX::Meis1cKO*. Statistical analysis was performed using two-tailed t-tests. p values are represented as **** for p < 0.0001 and N.S. for p > 0.05.

**Figure S12.**
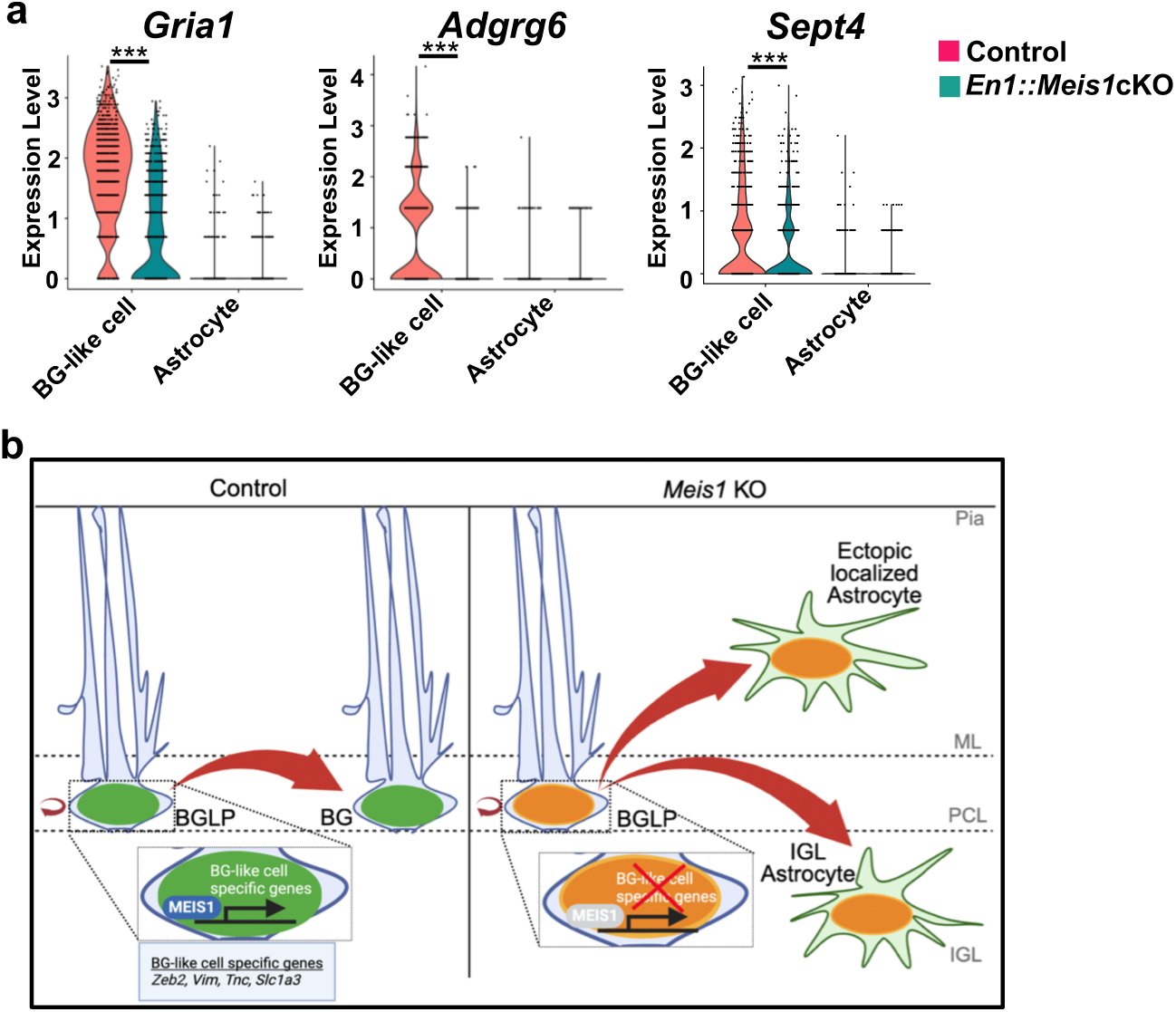
Supplemental data for Figure 5 (a) Violin plots showing the expression levels of BG-like cell–specific gene *Gria1, Adgrg6* and *Sept4* in scRNA-seq data. Expression values were compared between control and *En1::Meis1cKO*, separately for BG-like cells and astrocytes. Statistical analysis was performed using Wilcoxon Rank Sum test. p values are represented as *** for p < 0.001. (b) Schematic summary of the role of MEIS1 in regulating astroglial lineage output. In control (left), MEIS1 maintains the expression of BG-like cell–specific genes (*Zeb2, Vim, Tnc, Slc1a3*) within BGLPs and BGs, thereby promoting the endowment of BG-specific characteristics. In contrast, in *Meis1*-deficient (right), the loss of MEIS1 impairs the expression of these BG-like cell–specific genes, resulting in the loss of specialized astrocytes (BGs) from the cerebellum.

**Supplementary Table S1.** Genes downregulated and upregulated in BG-like cell subclusters of *En1::Meis1cKO* Genes showing significantly decreased or increased expression in BG-like cell subclusters of *En1::Meis1cKO* mice compared with controls. BG-like cells were defined based on subclusters 1 and 2 in the scRNA-seq dataset (Figure 1e). A total of 696 genes with significantly decreased expression and 420 genes with significantly increased expression in *En1::Meis1cKO* were identified and are listed in this table.

**Supplementary Table S2.** MEIS1 potential target genes List of MEIS1 potential target genes identified by integrating scRNA-seq and ChIP-Atlas analyses. According to ChIP-Atlas, a data-mining platform for epigenomic landscapes based on integrated ChIP-seq datasets (Oki et al., 2018; https://chip-atlas.org), 9,599 gene loci show MEIS1 binding in at least one cell type or tissue. Among the 696 genes downregulated in BG-like cell subclusters of *En1::Meis1cKO* mice, 355 genes overlapped with MEIS1-binding loci identified in ChIP-Atlas (Figure 5a). These overlapping genes are referred to as MEIS1 potential target genes in this study.

**Supplementary Table S3.** Primer sequences used for qPCR analysis List of primer pairs used for quantitative PCR (qPCR) analysis in this study. Primers were designed for the following genes: *Gapdh*, *Meis1*, *Sox9*, *Sox2*, *S100b*, *Vim*, *Zeb2*, *Gdf10*, *Tnc*, *Gria1*, *Septin4*, *Adgrg6*, *Glast*, *Slc6a11*, and *Mlc1*.

